# A genome compendium reveals diverse metabolic adaptations of Antarctic soil microorganisms

**DOI:** 10.1101/2020.08.06.239558

**Authors:** Maximiliano Ortiz, Pok Man Leung, Guy Shelley, Marc W. Van Goethem, Sean K. Bay, Karen Jordaan, Surendra Vikram, Ian D. Hogg, Thulani P. Makhalanyane, Steven L. Chown, Rhys Grinter, Don A. Cowan, Chris Greening

**Affiliations:** Centre for Microbial Ecology and Genomics, Department of Biochemistry, Genetics and Microbiology, University of Pretoria, Hatfield, Pretoria 0002, South Africa; Department of Microbiology, Monash Biomedicine Discovery Institute, Clayton, VIC 3800, Australia; School of Biological Sciences, Monash University, Clayton, VIC 3800, Australia; Environmental Genomics and Systems Biology Division, Lawrence Berkeley National Laboratory, Berkeley, California, USA; Departamento de Genética Molecular y Microbiología, Facultad de Ciencias Biológicas, Pontificia Universidad Católica de Chile, Alameda 340, Santiago; Securing Antarctica’s Environmental Future, School of Biological Sciences, Monash University, Clayton, VIC 3800, Australia; School of Science, University of Waikato, Hamilton 3240, New Zealand; Polar Knowledge Canada, Canadian High Arctic Research Station, Cambridge Bay, NU X0B 0C0, Canada

## Abstract

A surprising diversity and abundance of microorganisms resides in the cold desert soils of Antarctica. The metabolic processes that sustain them, however, are poorly understood. In this study, we used metagenomic and biogeochemical approaches to study the microbial communities in 16 physicochemically diverse mountainous and glacial soils from remote sites in South Victoria Land, north of the Mackay Glacier. We assembled 451 metagenome-assembled genomes from 18 bacterial and archaeal phyla, constituting the largest resource of Antarctic soil microbial genomes to date. The most abundant and prevalent microorganisms are metabolically versatile aerobes that use atmospheric hydrogen and carbon monoxide to meet energy, carbon, and, through metabolic water production, hydration needs. Phylogenetic analysis and structural modelling infer that bacteria from nine phyla can scavenge atmospheric hydrogen using a previously unreported enzyme family, the group 1l [NiFe]-hydrogenases. Consistently, gas chromatography measurements confirmed most soils rapidly consume atmospheric hydrogen and carbon monoxide, and provide the first experimental evidence of methane oxidation in non-maritime Antarctica. We also recovered genomes of microorganisms capable of oxidizing other inorganic compounds, including nitrogen, sulfur, and iron compounds, as well as harvesting solar energy via photosystems and novel microbial rhodopsins. Bacterial lineages defined by symbiotic lifestyles, including Patescibacteria, Chlamydiae, and predatory Bdellovibrionota, were also surprisingly abundant. We conclude that the dominant microorganisms in Antarctic soils adopt mixotrophic strategies for energy and sometimes carbon acquisition, though they co-exist with diverse bacteria and archaea that adopt more specialist lifestyles. These unprecedented insights and associated genome compendium will inform efforts to protect biodiversity in this continent.

## Introduction

Continental Antarctica is a relatively pristine but oligotrophic wilderness ^1^. Terrestrial life on the continent is adapted to extremely low temperatures, low water bioavailability, highly limited organic carbon and nitrogen, salt accumulation and seasonal light/dark periodicity ^2–4^. These cumulative pressures exclude most macroscopic fauna and flora, and instead microorganisms constitute most of the continent’s biodiversity and biomass ^5^. While historical observational surveys indicated that few microorganisms existed in terrestrial Antarctica, subsequent molecular studies have uncovered rich and abundant microbial communities, especially in the continent’s ice-free regions ^6–10^. Antarctic soil communities are comparable to mesophilic soils at the phylum level, with Actinobacteriota, Acidobacteriota, Chloroflexota and Proteobacteria often predominant ^2,8,9,11,12^. These communities are highly specialised at lower taxonomic levels ^7,8^, however, and have unique functional traits ^11,13^. Complementary culture-based studies have also isolated a growing number of taxa from the continent, although from relatively few phyla ^14–17^. Most community members are assumed to be extremely slow-growing or adopt dormant states to adapt to the physicochemical conditions of the continent ^18^. In turn, the formation of a microbial ‘seed bank’ may provide a means to maintain biodiversity ^19,20^.

An enduring question is what metabolic strategies enable soil microorganisms to meet energy and carbon needs on this continent ^2^. Even in dormant states, cells still require a net energy input to maintain cellular integrity, repair damaged macromolecules, and generate a basal membrane potential ^21,22^. Conventionally it was thought that Cyanobacteria and microalgae are the major primary producers in Antarctic soils and that they produce the organic carbon to sustain organoheterotrophic bacteria ^2,11^. However, oxygenic photoautotrophs are typically in low abundance (<1% of total bacterial community) outside lithic niches ^11,23^ and hence are unlikely to produce sufficient organic carbon to sustain the energy and carbon needs of the dominant community members. More recently, some Antarctic soil bacteria were shown to conserve energy and acquire carbon independently of photoautotrophs ^12^. Genome-centric metagenomic studies have revealed that bacteria from several phyla, including Actinobacteriota, consume molecular hydrogen (H_2_) and carbon monoxide (CO) from the atmosphere. By liberating electrons from these ubiquitous and diffusible trace gases, these bacteria sustain aerobic respiration and fix carbon even when preferred organic substrates are limiting ^12,24^. However, given the relatively few metagenome-assembled genomes (MAGs) recovered (21) and limited geographical scope of this previous study ^12^, it is unknown whether trace gas oxidation is a widespread strategy among Antarctic bacteria. Several molecular and biogeochemical studies have detected signatures of carbon fixation through the Calvin-Benson-Bassham (CBB) cycle within the continent, though it is unclear whether this originates through activities of photoautotrophs or lithoautotrophs ^12,13,25–27^. Molecular evidence also suggests that some Antarctic soil bacteria can also conserve energy through other means, including methanotrophy, nitrification, and rhodopsin-based light harvesting ^12,13,16,28–30^.

Here we build on these initial findings to develop a holistic genome-resolved understanding of the metabolic capabilities of Antarctic soil microorganisms. We profiled 16 soils with distinct physicochemical properties from the Mackay Glacier region, a cold hyper-arid ice-free region to the north of the McMurdo Dry Valleys that comprises approximately 15% (∼4,800 km^2^) of the ice-free regions on the continent. Soil microbial communities in this region are adapted to average annual temperatures of −20°C and annual precipitation below 50 mm ^31,32^, as well as profound limitation for organic carbon (∼0.1%) and nitrogen (∼0.02%) ^33^. Through deep metagenomic sequencing, we generated a resource of 451 metagenome-assembled genomes, covering all major microbial lineages in the region. We confirmed that the most abundant bacteria in the region are mixotrophs that scavenge atmospheric trace gases, and substantiated these findings with biogeochemical assays confirming rapid gas consumption and phylogenetic analyses revealing a novel hydrogenase family. These findings lend strong support to the recent hypothesis that survival in desert soils depends on continual harvesting of alternative energy sources ^18^. Nevertheless, these metabolically versatile bacteria co-exist with microorganisms that adopt a wide range of other nutritional and ecological strategies, including apparent obligate parasites and predators. Altogether, Antarctic soils appear to harbour much more compositionally rich and functionally complex microbial life than previously assumed.

## Results

### Genome-resolved metagenomics reveals phylogenetically diverse bacteria co-exist across the Mackay Glacier region

We analyzed surface soils from sixteen glacial and mountainous sites sampled across the Mackay Glacier region of South Victoria Land. Physicochemical analysis confirmed that the soils varied in key properties (e.g. pH, salinity, micronutrients, texture), but in common with previously characterized soils from continental Antarctic regions^8,34,35^, all had exceptionally low organic carbon content (0.02 – 0.25%) (**Table S1**). These soils nevertheless supported moderately abundant bacterial and archaeal communities (1.7 × 10^6^ to 2.7 × 10^7^ 16S rRNA gene copies per gram soil wet weight) (**Figure 1a**). Based on high-resolution 16S rRNA amplicon sequencing ^36^ (**Figure S1a & S1b**), observed richness (832 ± 258) and Shannon index (5.27 ± 0.31) were high in most samples, implying diverse community members co-exist in these soils (**Figure 1c; Figure S1d**). Beta diversity analysis confirmed microbial communities diverge between sampled regions and with geographic distance (**Figure 1d; Figure S1e**).

**Figure 1.**
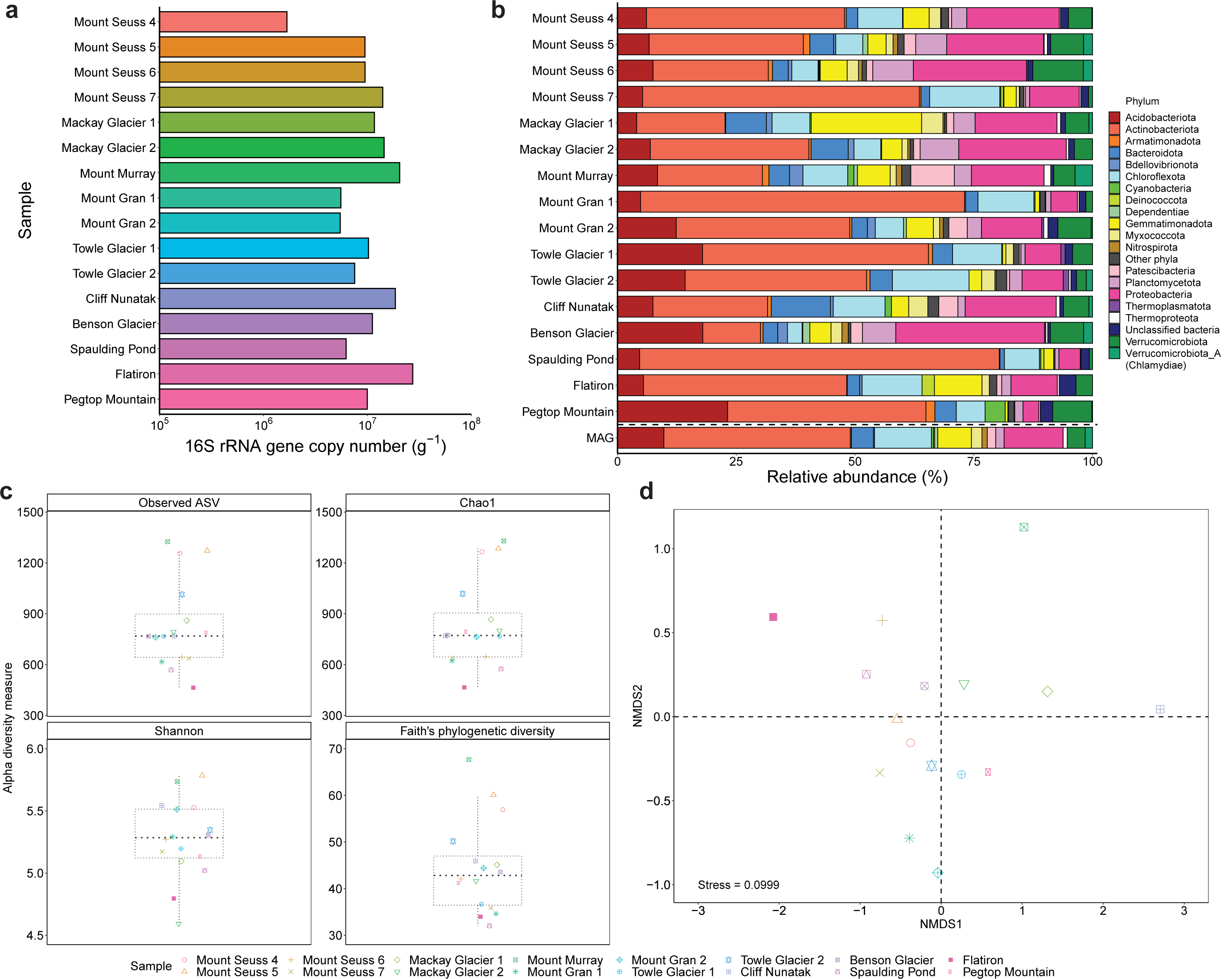
Abundance, composition, and diversity of the microbial communities from the Mackay Glacier region. **(a)** Boxplot showing the estimated abundance of bacterial and archaeal taxa, based on 16S rRNA copy number determined by quantitative PCR. **(b)** Stacked bar chart showing phylum-level community composition based on metagenomic reads of the single-copy marker gene *rplP* and metagenome-assembled genomes. Bacterial and archaeal taxonomy is based on Genome taxonomy database (GTDB) release 05-RS95. Phyla with less than 1% abundance in the sample were grouped to “Other phyla”. **(c)** Boxplot showing alpha diversity (Observed richness, Chao1, Shannon, Faith’s phylogenetic diversity) of microbial communities based on 16S rRNA gene amplicon sequence variants. **(d)** Beta diversity of rarefied 16S rRNA gene amplicon sequencing data based on Bray-Curtis dissimilarity and visualised by a non-metric multidimensional scaling ordination (NMDS) plot.

To determine the community composition of the samples, we retrieved and classified shotgun metagenomic reads of the universal single-copy ribosomal protein gene *rplP* (**Table S2**). The dominant community members were from bacterial phyla known to predominate in soil ecosystems ^37,38^. Actinobacteriota, Proteobacteria, Acidobacteriota, Chloroflexota, Gemmatimonadota, Verrucomicrobiota and Bacteroidota were particularly abundant (**Figure 1b**), in agreement with other Antarctic surveys ^2,18^. Cyanobacteria were scarce in most soils except for Pegtop Mountain and Cliff Nunatak, accounting for an average of 0.50% in the soil communities. Likewise, Archaea were minor members of this ecosystem (av. 0.88%) and mainly comprised the ammonia-oxidizing order Nitrososphaerales (**Figure 1b**). More surprisingly, bacterial phyla that predominantly adopt a predatory (Bdellovibrionota) ^39^, intracellular parasitic (Dependentiae and Verrucomicrobiota_A / Chlamydiae) ^40,41^ or obligately symbiotic (Patescibacteria) ^42,43^ lifestyle were prevalent and sometimes highly abundant, for example together comprising 17% of the community at Mount Murray. This suggests that a range of symbiotic interactions occur in these communities. These dominant and rare phyla were also detected by 16S rRNA gene sequencing **(Figure S1c; Table S3**).

These inferences on the composition and metabolic capabilities of the microbial communities were supported by genome-resolved analysis. From the 99.5 gigabases of sequencing data (**Table S4**), we reconstructed a non-redundant set of 101 high-quality and 350 medium-quality ^44^ metagenome-assembled genomes (MAGs). The recovered genomes span 18 different phyla, the relative composition of which reflects the community structure patterns observed in the *rplP* and 16S rRNA analysis (**Figure 1b**). In turn, they capture all major microbial lineages (present at >1% relative abundance across all samples) and map to an average of 26% of reads in each metagenome (**Table S5**). To the best of our knowledge, this represents the largest sequencing effort and most extensive genomic resource reported from terrestrial Antarctica to date.

### Most abundant lineages encode enzymes supporting trace gas oxidation, including a novel family of [NiFe]-hydrogenases

We sought to understand which metabolic strategies support the numerous bacteria in these hyper-oligotrophic soils. We profiled the distribution and affiliation of 52 conserved marker genes representing different energy conservation and carbon acquisition pathways in both the metagenomic short reads (**Table S6**) and MAGs (**Table S5**). In line with expectations, almost all community members encoded genes for aerobic organotrophic respiration (CoxA, NuoF, SdhA, AtpA) (**Figure 2**), whereas capacity for anaerobic respiration and fermentation was low (**Figure S2**). In addition to formate dehydrogenase, the other most abundant markers were the catalytic subunits of [NiFe]-hydrogenases (present in average of 90% community members), form I carbon monoxide dehydrogenases (32%), and RuBisCO (27%) (**Figure 2**). Phylogenetic analysis revealed that most binned sequences of these enzymes were most closely related to clades that support atmospheric H_2_ oxidation ^45–49^ (**Figure 3a**), atmospheric CO oxidation ^12,50–53^ (**Figure S3**), and chemosynthetic CO_2_ fixation ^12,54–56^ (**Figure S4**). Recent pure culture studies have shown that energy liberated by atmospheric H_2_ and CO oxidation supports bacterial persistence during carbon starvation and, in some cases, mixotrophic growth ^52,57–62^. Thus, the ability of bacteria to harvest these trace gases may confer a major selective advantage in the carbon-depleted soils of Antarctica. Moreover, in extension of findings made in the Windmill Islands region ^12^, over a quarter of the community may fix carbon via the CBB cycle, providing a mean to generate biomass independently of photoautotrophy.

**Figure 2.**
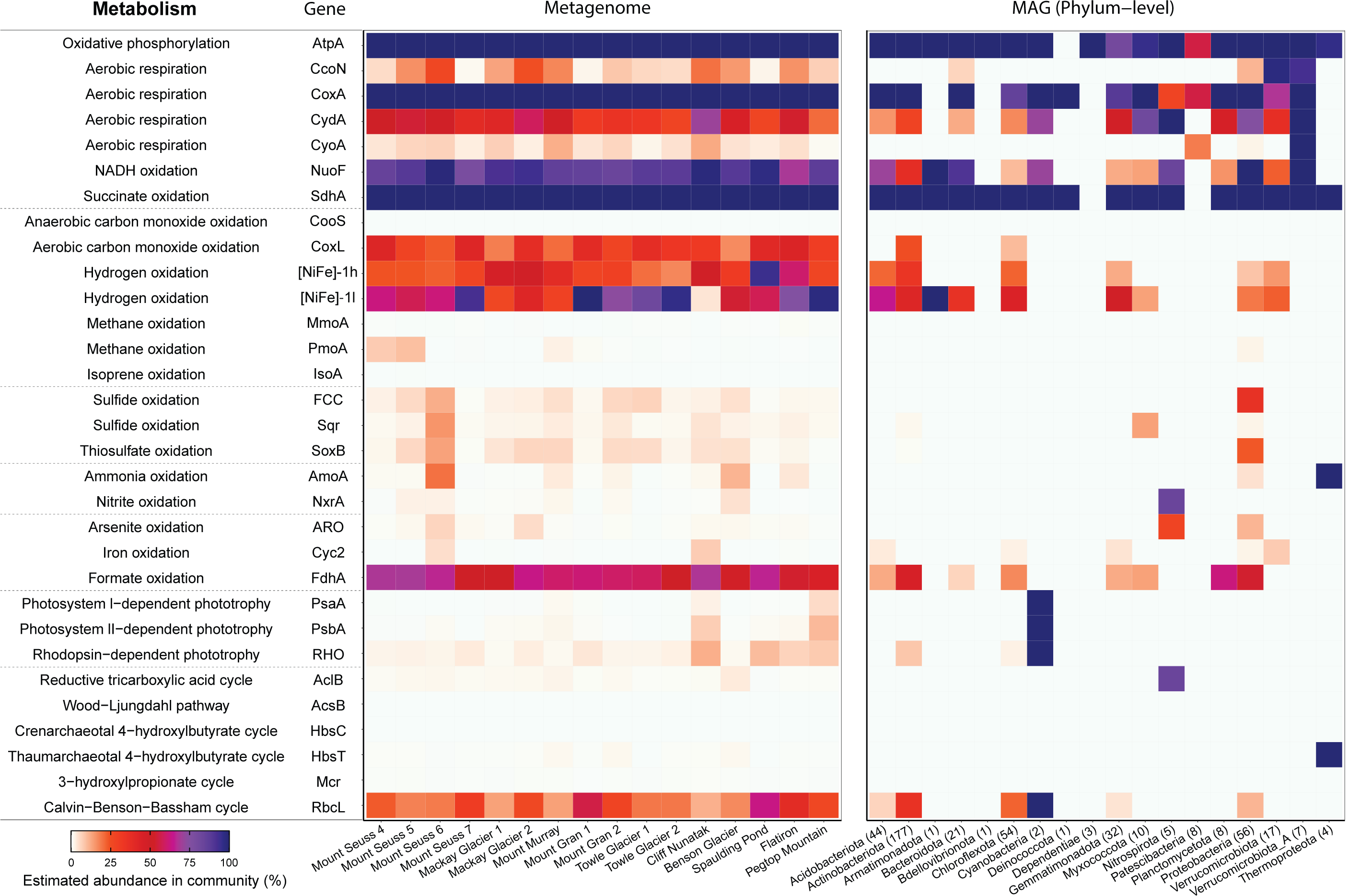
Metabolic potential of the microbial communities to use inorganic compounds, organic compounds, and light for energy and carbon acquisition. Homology-based searches were used to identify signature genes encoding enzymes associated with (from top to bottom): oxidative phosphorylation, trace gas oxidation, sulfur compound oxidation, nitrification, other oxidative processes, photosynthesis, and carbon fixation. The left heatmap shows the percentage of total community members predicted to encode each signature metabolic gene. To infer abundance, read counts were normalized to gene length and the abundance of single-copy marker genes. The right heatmap shows the presence of these genes across the 451 metagenome-assembled genomes spanning 18 phyla. Abundance was normalized by predicted MAG completeness.

**Figure 3.**
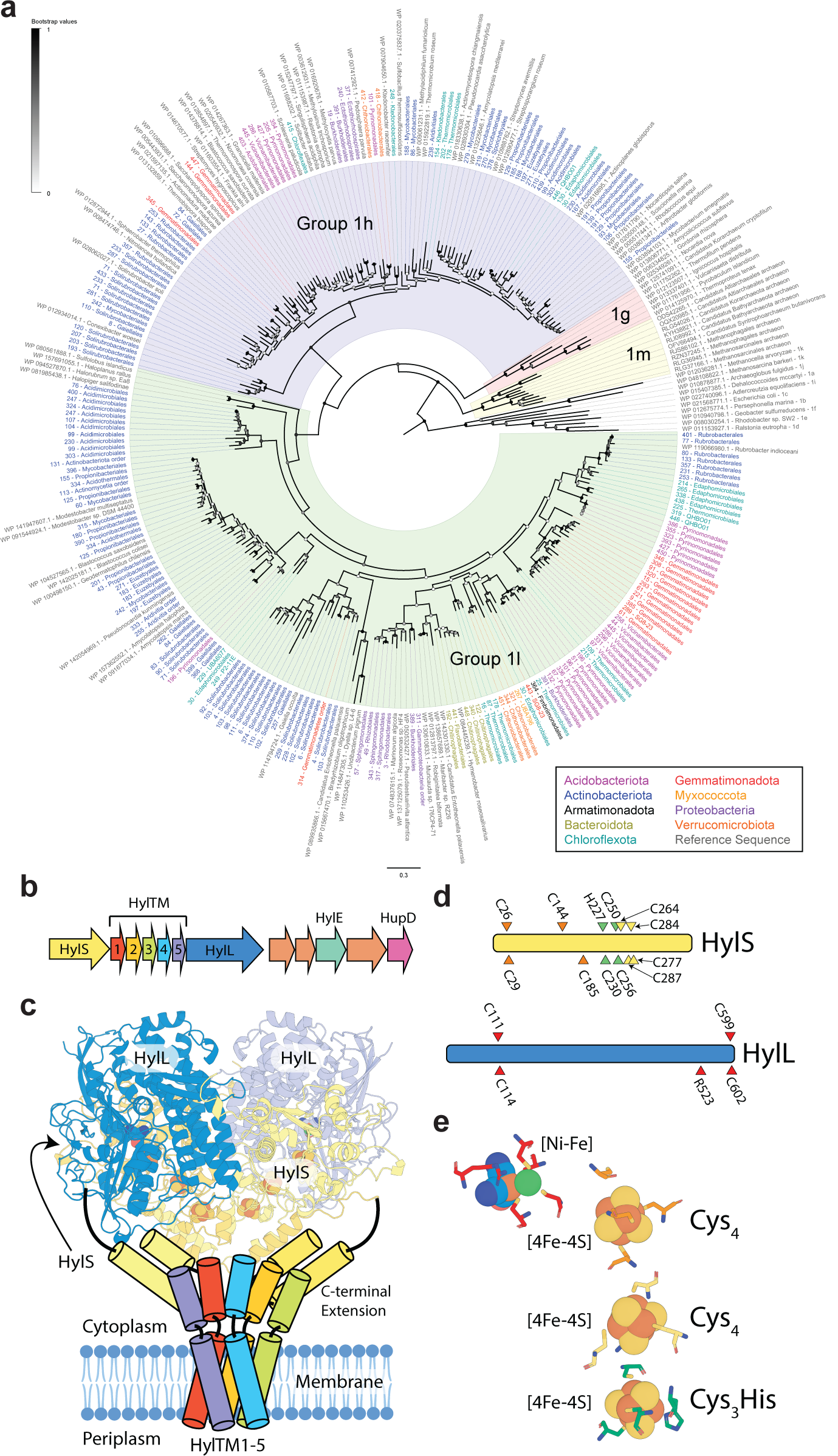
Identification of the novel group 1l family of [NiFe] hydrogenases widespread in the Antarctic soil bacterial communities. **(a)** Maximum-likelihood phylogenetic tree showing the sequence divergence of group 1 [NiFe] hydrogenases identified in MAGs from this study. Amino acid sequences retrieved from the reconstructed genomes were aligned against reference sequences (bootstrapped with 50 replicates). Branches of group 1 [NiFe] hydrogenases are shaded according to the subgroup classification and tips are colored based on phylum-level affiliation of the sequence. All sequences from MAGs of the Mackay Glacier region clustered with either the well-characterized group 1h [NiFe]-hydrogenases or the previously unreported group 1l [NiFe]-hydrogenases. **(b)** Representative genetic organization of group 1l [NiFe] hydrogenase gene cluster derived from the Antarctic bacterium *Hymenobacter roseosalivarius*. This shows the predicted open reading frames for the large (HylL) and small (HylS) hydrogenase subunits, the five interposing short predicted transmembrane proteins (HylTM1-5), a predicted electron-relaying Rieske-type protein (HylE), and a maturation endopeptidase (HupD). Conserved open reading frames with no predicted function are shown but not labelled. **(c)** Three-dimensional model of the group 1l [NiFe] hydrogenase. This shows a structural homology model of a heterotetramer of HylL and HylS subunits as a ribbon representation and a cartoon of a speculative complex between the hydrogenase and genetically associated HylTM proteins. **(d)** The location of conserved residues coordinating the [NiFe]-centre of the HylL subunit and [FeS] clusters of the HylS subunit of the group 1l [NiFe] hydrogenase. **(e)** Putative location of [FeS] clusters and [NiFe] centre (spheres) in one half of the group 1l [NiFe] hydrogenase tetramer, with conserved coordinating residues (sticks) color coded as in panel C.

Genes for trace gas oxidation were present in the most abundant and widespread community members. Uptake hydrogenases were encoded by MAGs affiliating with nine bacterial phyla (**Figure 2 & 3a**), including the seven dominant soil phyla (**Figure 1**), whereas CO dehydrogenases were confined to Actinobacteriota and Chloroflexota (**Figure S3**). Indeed, 17 of the 20 most abundant Actinobacteriota and Chloroflexota MAGs encoded one or both enzymes (**Table S5**). Remarkably, the CBB pathway (**Figure S4; Table S7**) frequently co-occurs with hydrogenases (64%) and CO dehydrogenase (25%) in MAGs (**Figure 2; Table S5**), potentially enabling hydrogenotrophic, carboxydotrophic or mixotrophic growth. This association was especially pronounced in the uncultivated classes Ellin6529 (Chloroflexota) and UBA4738 (Actinobacteriota) (**Table S6**), which respectively comprise an average of 5.1% and 0.9% (maximum of 12.3% and 2.4%) of the communities across the region (**Table S2**). These classes are predicted to couple atmospheric H_2_ and CO oxidation to fix carbon via their respective type IC and IE RuBisCO enzymes (**Figure S4; Table S7**). These traits in turn may contribute to their unexpectedly high relative abundance in Antarctica as well as other oligotrophic soils ^15,63–66^. Indeed, given their abundance in the community and genetic potential for atmospheric chemosynthesis ^12,24^, we hypothesize that both classes are major Antarctic primary producers. We propose replacing the placeholder names UBA4738 with *Candidatus* Aridivitia (arid Actinobacteriota class; based on high-quality type MAG MGR_bin238, ‘*Candidatus* Aridivita willemsiae’) and Ellin6529 with *Candidatus* Edaphomicrobia (edaphic Chloroflexota class; based on high-quality type MAG MGR_130 ‘*Candidatus* Edaphomicrobium janssenii’) (**Etymological Information**), as per recent taxonomic recommendations ^67,68^.

Most microorganisms in the Mackay Glacier region encoded a novel hydrogenase family (**Figure 2**). We generated a maximum-likelihood tree of the conserved catalytic subunits of group 1 [NiFe] hydrogenases using amino acid sequences retrieved from 176 MAGs. All hydrogenase sequences form two major and tremendously diverse lineages that share less than 40% sequence identity with each other and were supported by robust bootstrapping (**Figure 3a**). One branch is associated with characterized group 1h [NiFe] hydrogenases from multiple bacterial isolates ^45–47,51,61^. The other forms a novel cluster, herein the group 1l [NiFe]-hydrogenase, which includes the previously unreported hydrogenases of McMurdo Dry Valleys isolate *Hymenobacter roseosalivarius* ^69^ and several other recently sequenced isolates. Group 1l is the prevailing hydrogenase family within the Mackay Glacier region, with an estimated abundance 2.3 times higher than group 1h (**Table S5**), and is encoded by all nine hydrogenase-bearing phyla and the two candidate classes. As elaborated in **Supplementary Note 1**, structural modelling shows that this enzyme shares common structural features with previously characterized group 1h [NiFe]-hydrogenase ^70,71^, but contains large sequence insertions and a key substitution in a residue ligating the proximal iron-sulfur cluster. Even more strikingly, the genes encoding this hydrogenase often have an unusual arrangement (**Figure S5; Table S7**), with five open reading frames predicted to encode small transmembrane proteins separating the small and large core structural subunits. On this basis, we predict that this enzyme is a *bona fide* high-affinity membrane-associated hydrogenase that relays electrons derived from atmospheric H_2_ through the respiratory chain. The broad distribution and predominance of this hydrogenase suggests it is the primary mediator of H_2_ oxidation in these soils. Moreover, given the strong positive correlation between this hydrogenase and RuBisCO based on the MAGs and metagenomic short reads (*R*^2^ = 0.68, *p* = 0.002) (**Figure S6; Table S9**), it is likely that electrons yielded by this enzyme support carbon fixation either through direct transfer or reverse electron flow.

### Trace gas consumption occurs at sufficient rates to meet energy needs and support hydration of Mackay Glacier region bacteria

Our metagenomic analyses suggest that the most abundant soil bacteria across the Mackay Glacier region conserve energy and fix carbon by oxidizing atmospheric H_2_ and CO. To test whether soil communities mediate these activities, we set up soil microcosms in which ambient air headspaces were amended with 10 parts per million (ppmv) of these gases and used high-sensitivity gas chromatography to measure their consumption over time. In line with predictions, H_2_ was oxidized by soils from all sixteen sites and all but three soils consumed CO (**Figure 4a**). Of these, all soils except Pegtop Mountain consumed H_2_ to below atmospheric concentrations (0.53 ppmv) ^72^ and ten soils consumed atmospheric CO (0.09 ppmv) ^73^ during the timecourse of our experiments (**Figure S7**). These sub-atmospheric thresholds confirm that these microbial communities can harvest energy from the atmosphere, a virtually unlimited source of diffusive and energy-rich reduced gases ^74,75^. The average rate of atmospheric H_2_ oxidation (135 pmol hr^-1^ g_soil ww_ ^-1^) was much faster than for atmospheric CO oxidation (0.60 pmol hr^-1^ g_soil ww_^-1^) (**Table S8**). This finding, together with the higher abundance of putative H_2_ oxidizers in the soil communities (**Figure 2**), suggests that atmospheric H_2_ is likely to be the predominant energy source sustaining these communities. As elaborated in **Supplementary Note 2**, considerable variations in bulk and normalized oxidation rates were measured for both gases, which was significantly correlated with several measured physicochemical variables (**Figure S6; Table S9**).

**Figure 4.**
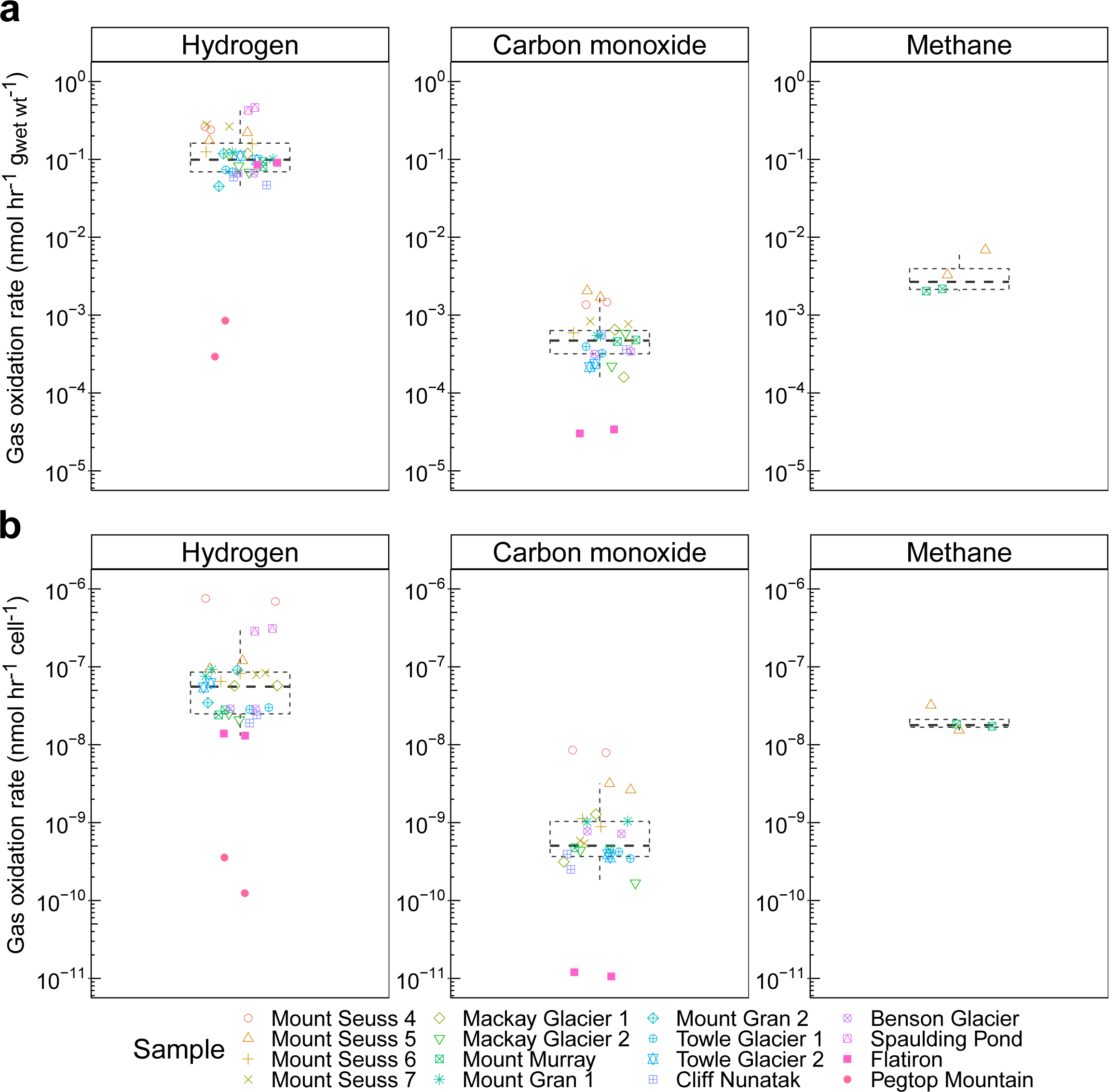
Rates of atmospheric trace gas oxidation by soils sampled from the Mackay Glacier region. Boxplots show rates of oxidation of atmospheric H_2_, CO, and CH_4_ for each soil in duplicate soil microcosms at 10°C, based on gas chromatography measurements. Only rates for samples with detectable gas oxidation are shown. **(a)** Atmospheric gas oxidation rate for each microcosm normalized to wet weight of soil. **(b)** Cell-specific reaction rates for each microcosm. These rates were calculated by dividing the estimated soil cell abundance and proportion of gas oxidizers based on quantitative qPCR and metagenome short read analysis (HhyL and HylL abundance for H_2_, CoxL abundance for CO, PmoA and MmoX abundance for CH_4_).

Cell-specific rates were calculated by normalizing bulk rates against soil microbial abundance and the proportion of trace gas oxidizers. Cell-specific atmospheric H_2_ oxidation rates were high (av. 1.1 × 10^−7^ nmol hr^-1^ cell^-1^) and approximately two orders of magnitude higher than those of CO (av. 1.3 × 10^−9^ nmol hr^-1^ cell^-1^) (**Figure 4b**). In line with our findings in the Windmill Islands region ^12^, this rate of atmospheric H_2_ consumption exceeds the theoretical maintenance requirements of trace gas oxidizers at the temperature tested (10°C) and is sufficient to support some growth ^76–78^. It should also be noted that metabolic water is the major end-product of the aerobic respiration of atmospheric H_2_ (2 H_2_ + O_2_ → 2 H_2_O). Given the reported cytosolic orientation of high-affinity hydrogenases and terminal oxidases ^62^, the water produced would be retained in the cytosol, including as a solvent for macromolecules. Thus, trace gas oxidation may be a simple, but hitherto overlooked, mechanism for microorganisms to stay hydrated in the hyper-arid deserts of Antarctica. Based on cell-specific rates of atmospheric H_2_ oxidation, a theoretical average of 1.1 million water molecules would be produced per cell each minute. For a cell with an expected 1 µm^3^ volume and 70% water content ^79,80^, such production rates would be sufficient to replace all cellular water over a 15-day period (**Table S8**). We therefore propose that the metabolic water continuously generated by trace gas oxidation is a quantitatively significant source of hydration in this environment with minimal precipitation ^32^.

### Metabolically constrained phototrophs, lithotrophs, and organotrophs co-exist with versatile mixotrophs in Antarctic soils

While the most abundant taxa in the Mackay Glacier ecotone appear to be versatile mixotrophs, the genome compendium revealed that these ecosystems also harbor diverse bacteria and archaea with specialist strategies for energy and carbon acquisition. Multiple chemolithoautotrophs were present, including those capable of oxidizing the trace amounts of ammonium, sulfur and iron detected in the soils (**Table S1**). Ammonium and nitrite oxidizers comprised an average of 2.9% and 1.0% of the communities, but together comprised 23% and 15% of the community in Mount Seuss 6 and Benson Glacier samples, respectively (**Figure 2; Table S6**). Phylogenetic analysis confirmed that Nitrososphaerales (archaea) and Burkholderiales (bacteria) were the dominant ammonium oxidizers (**Figure S8**), in line with previous reports for McMurdo Dry Valley soils ^28^, whereas Nitrospirota were the main nitrite oxidizers (**Figure S9**). These nitrifiers also respectively encoded the signature enzymes to fix carbon through the archaeal 4-hydroxybutyrate cycle (**Figure S10**), proteobacterial CBB cycle (**Figure S4**), and nitrospiral reverse tricarboxylic acid cycle (**Figure S11**), suggesting that multiple chemosynthetic primary production strategies sustain biodiversity in these oligotrophic soils. The marker genes for sulfide and thiosulfate oxidation (Sqr, FCC, SoxB) were each encoded by 1 - 4% of community members in most soils (**Figure 2; Table S5**), including multiple Burkholderiales MAGs and several other lineages (**Figure S12, S13, S14**). The genes to oxidize ferrous iron via the *c*-type cytochrome Cyc2 were widespread in Mount Seuss 6 (4.7%) and Cliff Nunatak samples (7.3%), and present in select MAGs from five major phyla (**Figure S15**). Thus, atmospheric and edaphic inorganic compounds alike are major energy sources for Antarctic soil communities, although their relative importance varies across the physicochemically diverse soils from the region.

Our metagenomic analysis suggests that light energy supports few photoautotrophs, but numerous photoheterotrophs, in the region. Reflecting cyanobacterial distributions across the region (**Figure 1b**), photosystems associated with oxygenic photosynthesis were encoded by few community members except in the Pegtop Mountain and Cliff Nunatak samples (**Figure 2**). Some photosystem II sequences affiliated with proteobacterial anoxygenic phototrophs were also detected (**Figure S16**). In contrast, energy-converting microbial rhodopsins were prevalent and abundant across the region (**Figure 2**). These light-powered proton pumps are well-characterized for their role in energy conservation in marine and freshwater ecosystems ^81–85^, though have been scarcely studied in desert environments ^86^. As outlined by our ‘continual energy harvesting hypothesis’, sunlight (in common with atmospheric trace gases) is a relatively dependable energy source and hence lineages that harvest it may have a selective advantage in energy-poor desert soils ^18^. In line with this theory, putative energy-converting rhodopsins were present in several of the most dominant orders of Actinobacteriota and Chloroflexota in these soils (**Table S5**). They were also present in both cyanobacterial MAGs, thereby providing a means for photoautotrophs to conserve energy when water for oxygenic photosynthesis is limiting (**Figure S17**). Phylogenetic analysis confirmed the binned and unbinned sequences fell into diverse clades (**Figure S17**), including two novel clades that were most closely related (<50% sequence identity) to the biochemically characterized energy-converting rhodopsins of halophilic archaea (bacteriorhodopsins) ^87^ and *Pantoea* species (pantorhodopsins) ^88^.

Twenty metagenome-assembled genomes were also recovered for the phyla known to adopt obligately symbiotic lifestyles, namely Patescibacteria, Chlamydiae, Dependentiae, and Bdellovibrionota (**Table S5**). All four phyla appear to be obligate organoheterotrophs that lack alternative pathways for energy conservation or carbon acquisition (**Figure 2**). Based on previous reports, all characterized Bdellovibrionota predate bacterial species ^39^, whereas Chlamydiae and Dependentiae are likely to be parasites of protist or arthropod species ^40,41,89^ such as populations of springtails (Collembola) identified within the same sampling area ^90^. Signature genes associated with the symbiotic lifestyles of each MAG were detected, for example host-targeted peptidoglycan metalloendopeptidases and self-protection proteins that Bdellovibrionota uses to invade cells of bacterial prey ^91,92^, as well as ankyrin repeat and WD40 repeat proteins implicated in modulation of eukaryotic hosts by Dependentiae ^41,89^ (**Table S5**). Also in line with an obligately symbiotic lifestyle, several lineages have ultra-small genomes when adjusted for completeness, namely the eight Patescibacteria MAGs (av. 1.3 Mbp), three Dependentiae MAGs (av. 1.8 Mbp), and a Rickettsiaceae MAG (1.3 Mbp) (**Table S5**), and are predicted to be auxotrophic for multiple amino acids. Building on the discovery of unexpected symbionts in Antarctic lakes ^93,94^, to our knowledge this is the first report that microbial parasitism is a major ecological strategy in terrestrial Antarctica. We also reveal oxic niches for phyla such as Patescibacteria that have, until now, primarily been studied in anoxic ecosystems ^42,95,96^.

Finally, we obtained genomic and biogeochemical evidence that atmospheric methane oxidation occurs on non-maritime Antarctic soils. Based on methane monooxygenase levels in short reads, aerobic methanotrophs are members of the rare biosphere in most of the sampled Antarctic soils, but are present in very high levels in three soils, including Mount Seuss 5 (9.4%) (**Figure 2; Table S5**). Concordantly, two of these soils oxidized methane at high cell-specific rates to sub-atmospheric levels during microcosm incubations (**Figure 4; Figure S7**). Genome-resolved analysis suggested that this activity is primarily mediated by a single bacterial species within the gammaproteobacterial order UBA7966, which encodes a particulate methane monooxygenase clustering with sequences from the atmospheric methane-oxidizing clade USCγ (**Figure S18**). While this bacterium has a restricted distribution, based on read mapping, it is among the most abundant single taxon across the entire region (**Table S5**). Thus, by adopting a relatively specialist lifestyle dependent on assimilating a widely available but catalytically demanding atmospheric substrate, this bacterium fills a distinct ecological niche. Importantly, although methanotroph genomes have previously been reported in Antarctic soils ^12,30^, this is the first experimental report that such bacteria are biogeochemically active.

## Conclusions

Altogether, these results demonstrate a remarkable diversity of both microbial lineages and metabolic strategies in the resource-poor soils of Antarctica. The most abundant and prevalent bacterial lineages in Antarctic soils appear to be free-living mixotrophs capable of meeting carbon, energy, and even hydration needs from atmospheric trace gases, i.e. ‘living on air’ ^97^. Several bacteria and archaea also achieve high abundances in specific soils through more specialist strategies, spanning atmospheric methanotrophy, oxygenic photosynthesis and lithoautotrophic growth on trace edaphic substrates. This environment in turn has selected for a range of as-yet-uncultivated bacterial lineages (e.g. *Ca*. Edaphomicrobia and *Ca*. Aridivitia) and previously unreported gene families (e.g. encoding group 1l [NiFe]-hydrogenases and potential microbial rhodopsins). Also, surprisingly, a significant minority of community members gain resources through parasitism or predation of microorganisms. Through this combination of strategies, both free-living and symbiotic microorganisms can achieve stable niches in a polyextreme environment.

Additionally, the wealth of metagenomic sequencing data and 451 draft genomes generated by this study provides a valuable resource for two major areas of endeavor. First, these datasets support fundamental research and potentially inform decisions to secure Antarctica’s environmental future, given forecasts of changing temperature and water availability ^98–100^. Thus, in line with one of the six priorities for Antarctic science ^101^, this resource will provide insights into how life has evolved and adapted on this microbially-dominated continent, and in turn may respond to climate changes. Secondly, these findings also contribute to considerations of what processes may sustain life on other cold, dry planets such as Mars. Antarctica has long been considered a potential analogue for life elsewhere in the solar system ^102^. Our work brings that picture into sharper resolution.

## Materials and Methods

### Soil physicochemical analysis

This study used mineral soils previously sampled from 16 glacier- or mountain-associated sites in the Mackay Glacier region, South Victoria Land, Antarctica during January 2015 as previously described ^33,35^. In brief, 50 g of surface soil (depth: 0 - 5 cm) at each location was collected from an approximately 1 m^2^ area and stored in sterile 50 ml polypropylene Falcon tubes (Grenier, Bio-One) aseptically. During storage and transportation to University of Pretoria, samples were kept at −80°C. They were later shipped to Monash University’s quarantine approved facilities for further experiments. Details of soil samples can be found in **Table S1**. Prior to physicochemical measurements, approximately 35 g of soil of individual sample was aliquoted. Soil aliquots were treated with gamma irradiation at 50 kGy (Steritech Pty Ltd Victoria, Australia) for compliance with Department of Agriculture, Water and the Environment’s quarantine good regulations. They were subsequently shipped to the Environmental Analysis Laboratory (EAL), Southern Cross University, Australia for physicochemical analyses in accordance with ISO/IEC 17025 standard procedures. Physicochemical parameters analysed included: basic soil colour and texture; pH and electrical conductivity (1:5 water); moisture content; total carbon, nitrogen, organic carbon, and organic matter; available calcium, magnesium, potassium, ammonium, nitrate, phosphate, sulfur; exchangeable sodium, potassium, calcium, magnesium, hydrogen, and aluminium; cation exchange capacity; Bray I, Bray II, and Cowell phosphorus; and available micronutrients zinc, manganese, iron, copper, boron, and silicon. These data are summarised in **Table S1**.

### Shotgun metagenome sequencing, assembly and binning

Community DNA for metagenomic sequencing was extracted from 0.5 g of soil using the FastDNA SPIN Kit for soil (MP Biomedicals) according to the manufacturer’s instructions. An extraction blank control was included. Metagenomic shotgun libraries were prepared using the Nextera XT DNA Sample Preparation Kit (Illumina Inc., San Diego, CA, USA) and subject to paired-end sequencing (2 × 150 bp) on an Illumina NextSeq500 platform at the Australian Centre for Ecogenomics (ACE), University of Queensland. Sequencing yielded 356,941,066 read pairs across the sixteen soil metagenomes and 556 read pairs for the negative control (**Table S4**), indicating a minimal level of contamination from DNA extraction and sequencing processes. Raw metagenomic sequences were subjected to quality filtering using the BBDuk function of the BBTools v38.80 (https://sourceforge.net/projects/bbmap/); contaminating adapters (k-mer size of 23 and hamming distance of 1), PhiX sequences (k-mer size of 31 and hamming distance of 1), and bases from 3’ ends with a Phred score below 20 were trimmed. After removing resultant reads with lengths shorter than 50 bp, 93% high-quality read pairs were retained for downstream analysis. Metagenomic reads from each sample were assembled individually with metaSPAdes v3.14.0 ^103^ and collectively with MEGAHIT v1.2.9 ^104^ (min k: 27, max k: 127, k step: 10). To generate corresponding coverage profiles for assembled contigs, short reads were mapped back using Bowtie2 v2.3.5 ^105^ with default parameters. Subsequently, genome binning was performed using CONCOCT v1.1.0 ^106^, MaxBin2 v2.2.7 ^107^, and MetaBAT2 v2.15 ^108^ on contigs with length over 2000 bp. Resulting bins from the same assembly were then dereplicated using DAS_Tool v1.1.2 ^109^. RefineM v0.0.25 ^110^ was used to remove spurious contigs with incongruent genomic and taxonomic properties. Applying a threshold average nucleotide identity of 99%, bins from different assemblies were consolidated to a non-redundant set of metagenome-assembled genomes (MAGs) using dRep v2.5.4 ^111^. Completeness and contamination of MAGs were assessed using CheckM v1.1.2 ^112^. In total, 101 high quality (completeness > 90% and contamination < 5%) and 350 medium quality (completeness > 50% and contamination < 10%) ^44^ MAGs from 18 phyla were recovered. Their corresponding taxonomy was assigned by GTDB-TK v1.3.0 ^113^ with reference to GTDB R05-RS95 ^114^. Open reading frames (ORFs) in MAGs were predicted using Prodigal v2.6.3 ^115^.

### Community analysis

Soil microbial community structures were determined by using both metagenomic and 16S rRNA gene amplicon sequencing. Community profiles in sequenced metagenomes were generated by mapping quality-filtered reads to the universal single copy ribosomal marker genes and clustering at 97% identity using SingleM v.0.12.1 (https://github.com/wwood/singlem). To align with the latest GTDB taxonomy at the time of submission (R05-RS95; release 2020/07), we generated a SingleM package for the single-copy ribosomal protein-encoding gene *rplP*. In brief, all *rplP* sequences from Archaea and Bacteria genomes in GTDB R05-RS95 (https://data.ace.uq.edu.au/public/gtdb/data/releases/release95/95.0/) were downloaded. GraftM v0.12.2 ^116^ was used to generate a phylogenetic package for the sequences which was then used to make a community classification package by SingleM v.0.12.1. For 16S rRNA gene amplicon sequencing, the DNeasy PowerSoil kit (Qiagen) was used to extract DNA from 0.4 g of soil sample as per manufacturer’s instructions. The quality and concentration of DNA extracted were determined using a Nanodrop spectrophotometer (ND-1000) and a Qubit Fluorometer. Quantitative PCR (qPCR) using a 96-well plate in a pre-heated LightCycler 480 Instrument II (Roche, Basel, Switzerland) was used to quantify the copy number of the 16S rRNA genes in the samples as previously described ^117^. For each sample, the V4 hypervariable region for 16S rRNA gene was amplified using the universal Earth Microbiome Project primer pairs F515 (Parada) ^118^ and R806 (Apprill) ^119^. Amplicons were sent to paired-end sequencing (2 × 300 bp) on an Illumina MiSeq platform at the Australian Centre for Ecogenomics (ACE), University of Queensland. BBDuk function of the BBTools v38.80 was used to trim adapter sequences and filter PhiX contaminants as described above. The sequences were further processed on the QIIME2 platform (release 2019/07) ^120^ to resolve amplicon sequence variants (ASVs) through the following steps: (i) striping amplicons primers using cutadapt plugin ^121^; (ii) merging paired-end reads using q2-vsearch plugin ^122^; (iii) quality filtering using a sliding window of four bases with an average Phred score 20; and (iv) de-noising and truncating sequences at 250 base pairs using deblur ^123^. A total of 657,975 reads remained in the dataset (min: 13248, max: 102382) (**Table S3**). For taxonomic assignment, ASVs were independently annotated with trained naïve Bayes classifiers of 16S rRNA reference databases Silva release 138 ^124^ and Greengenes 13.8 ^125^ (**Table S3**). Multiple sequence alignment of the sequences and subsequent phylogenetic tree building were performed using MAFFT ^126^ and FastTree ^127^, respectively, implemented in QIIME2. We then used R packages phyloseq ^128^, picante ^129^, vegan ^130^, betapart ^131^ and ggplot2 ^132^ for downstream statistical analysis and visualizations. Alpha diversity including observed richness, Chao1, Shannon index, and Faith’s phylogenetic diversity were computed using estimate_richness function in phyloseq and pd function in picante. For beta diversity analysis, all samples were rarefied at the lowest sample sequencing depth, i.e. 13248 sequences per sample and rarefaction plots before and after rarefaction were shown in **Figure S1a-b**. Bray-Curtis dissimilarity was calculated and visualized using a non-metric multidimensional scaling ordination (NMDS) plot. To examine community turnover in relations to increasing geographic separation, a distance decay relationship of beta diversity (Bray-Curtis dissimilarity) against pairwise geographic distance was computed using the decay.model function fitted with a negative exponential law function in betapart. A *p* value was calculated using the same function with 999 permutations (**Table S3**).

### Functional analysis

To estimate the metabolic capability of the soil communities, metagenomes and derived genomes were searched against custom protein databases of representative metabolic marker genes using DIAMOND v.0.9.31 (query cover > 80%) ^133^. Searches were carried out using all quality-filtered unassembled reads with lengths over 140 bp and the ORFs of the 451 MAGs. These genes are involved in sulfur cycling (AsrA, FCC, Sqr, DsrA, Sor, SoxB), nitrogen cycling (AmoA, HzsA, NifH, NarG, NapA, NirS, NirK, NrfA, NosZ, NxrA, NorB), iron cycling (Cyc2, MtrB, OmcB), reductive dehalogenation (RdhA), phototrophy (PsaA, PsbA, energy-converting microbial rhodopsin), methane cycling (McrA, MmoA, PmoA), hydrogen cycling (catalytic subunit of [NiFe]-hydrogenases, catalytic domain of [FeFe]-hydrogenases, and Fe-hydrogenases), isoprene oxidation (IsoA), carbon monoxide oxidation (CoxL, CooS), succinate oxidation (SdhA), fumarate reduction (FrdA), and carbon fixation (RbcL, AcsB, AclB, Mcr, HbsT, HbsC) ^48,52,134^. Results were filtered based on an identity threshold of 50%, except for group 4 [NiFe]-hydrogenases, [FeFe]-hydrogenases, CoxL, AmoA, and NxrA (all 60%), PsaA (80%), PsbA and IsoA (70%), and HbsT (75%). Subgroup classification of reads was based on the closest match to the sequences in databases. To search for the presence of an additional set of genes involved in oxidative phosphorylation (AtpA), NADH oxidation (NuoF), aerobic respiration (CoxA, CcoN, CyoA, CydA), formate oxidation (FdhA), arsenic cycling (ARO, ArsC), and selenium cycling (YgfK), corresponding in-house databases were generated for this study. All archaeal and bacterial non-redundant proteins were retrieved from NCBI Refseq protein database release 99 ^135^, which were then screened by hidden Markov models (HMM) ^136^, with search cutoff scores as described previously ^137^. Resulting hits were manually inspected to remove false positives and genes with lengths that deviated more than 20% from the average were discarded. The search of these genes in unassembled reads and ORFs of MAGs was carried out using the DIAMOND blastp algorithm with a minimum percentage identity of 60% (NuoF), 70% (AtpA, ARO, YgfK) or 50% (all other databases). Read counts for each gene were normalized to reads per kilobase per million (RPKM) by dividing the actual read count by the total number of reads (in millions) and then dividing by the gene length (in kilobases). In order to estimate the gene abundance in the microbial community, high-quality unassembled reads were also screened for the 14 universal single copy ribosomal marker genes used in SingleM v.0.12.1 and PhyloSift ^138^ by DIAMOND (query cover > 80%, bitscore > 40) and normalized as above. Subsequently, the average gene copy number of a gene in the community was calculated by dividing the read count for the gene (in RPKM) by the mean of the read counts of the 14 universal single copy ribosomal marker genes (in RPKM).

### Phylogenetic analysis

Maximum-likelihood phylogenetic trees were constructed to verify the presence and visualise the evolutionary history of key metabolic genes in the metagenome-assembled genomes and assembled unbinned reads. Trees were constructed using the amino acid sequences for subunits of ten enzymes involved in energy acquisition: group 1 [NiFe]-hydrogenase (HhyL, HylL); form I carbon monoxide dehydrogenase (CoxL), particulate methane monooxygenase (PmoA), ammonia monooxygenase (AmoA), nitrite oxidoreductase (NxrA), sulfide-quinone oxidoreductase (Sqr), flavocytochrome *c* sulfide dehydrogenase (FCC), thiosulfohydrolase (SoxB), iron-oxidizing *c*-type cytochrome (Cyc2), photosystem II (PsbA), and energy-converting rhodopsins. Trees were also constructed of the amino acid sequences for subunits of three enzymes involved in carbon fixation: ribulose 1,5-bisphosphate carboxylase/oxygenase (RuBisCO; RbcL), thaumarchaeotal 4-hydroxybutyrate synthase (HbsT), and ATP-citrate lyase (AclB). In all cases, protein sequences retrieved from the MAGs or assembled metagenome sequences by homology-based searches were aligned against a subset of reference sequences from the custom protein databases using ClustalW ^139^ in MEGA X ^140^. Evolutionary relationships were visualized by constructing maximum-likelihood phylogenetic trees; specifically, initial trees for the heuristic search were obtained automatically by applying Neighbour-Join and BioNJ algorithms to a matrix of pairwise distances estimated using a JTT model, and then selecting the topology with superior log likelihood value. All residues were used and trees were bootstrapped with 50 replicates. To characterise the genetic context of [NiFe] hydrogenases and ribulose-1,5-bisphosphate carboxylase / oxygenase (RuBisCO) from the MAGs, up to 10 genes upstream and downstream of the catalytic subunits were retrieved. These flanking genes were annotated against Pfam protein family database v33.1 ^141^ using PfamScan v1.6 ^142^ and NCBI Refseq protein database release 99 ^135^ using DIAMOND^133^ blastp algorithm (default parameters). Alignments with the highest score were retained and are summarised in **Table S7**. The R package gggenes (https://github.com/wilkox/gggenes) was used to construct gene arrangement diagrams.

### Hydrogenase sequence analysis and homology modelling

The amino acid sequence for the large (HylL; GBID = SMB94678) and small subunits (HylS GBID = SMB94698) of the group 1l [NiFe]-hydrogenase from *H. roseosalivarius* were inputted into the Phyre2 webserver using default parameters ^143^. The highest confidence output model for both subunits was derived from the structure of the group 1h [NiFe]-hydrogenase (HhyLS) from *Cupriavidus necator* H16 (PDB ID = 5AA5) ^71^. The structure of the group 1l [NiFe]-hydrogenase tetramer was assembled using Pymol, based on the tetrameric structure of the *C. necator* group 1h [NiFe]-hydrogenase for further analysis. To identify transmembrane helix presence, position and topology in the HylTM proteins associated with group 1I [NiFe]-hydrogenases, the amino acid sequences from *H. roseosalivarius* were inputted into the TMHHM 2.0 webserver ^144^.

### Gas chromatography assays

Soil microcosms were used to determine the capacity of soil microbial communities to oxidize H_2_, CO, and CH_4_ by gas chromatography. For each of the 16 Mackay Glacier region samples in technical duplicate, 2 g of soil was placed in a 120 ml serum vial and incubated at 10°C. The ambient air headspace was amended with H_2_, CO, and CH_4_ (via a mixed gas cylinder containing 0.1 % v/v H_2_, CO, and CH_4_ each in N_2_, BOC Australia) to give starting mixing ratios of approximately 10 parts per million (ppmv) for each gas. At each time interval, 2 ml of headspace gas was sampled using a gas-tight syringe and stored in sealed a 3 ml glass exetainer that had been flushed with ultra-high purity N_2_ (99.999% pure, BOC Australia) prior to measurement. A VICI gas chromatographic machine with a pulsed discharge helium ionization detector (model TGA-6791-W-4U-2, Valco Instruments Company Inc.) and an autosampler was used to measure gas concentrations as previously described ^51^. The machine was calibrated against ultra-pure H_2_, CO and CH_4_ standards down to the limit of quantification (H_2_: 20 ppbv; CO: 9 ppbv; CH_4_: 500 ppbv). Calibration mixed gas (10.20 ppmv of H_2_, 10.10 ppmv of CH_4_, 9.95 ppmv of CO in N_2_, Air Liquide Australia) and pressurized air (Air Liquide Australia) with known trace gas concentrations were used as internal reference standards. Four pooled heat-killed soils (2 g of pooled soil; treated at 121°C, 15 p.s.i. for 60 mins) were prepared as negative controls. For kinetic analysis, measurement time points with individual gas concentration over 0.4 ppmv were used. First order reaction rate constants were calculated by fitting an exponential model as determined by the lowest overall Akaike information criterion value when compared to a linear model. Actual reaction rate constants of the sample were obtained by correcting against means of negative controls and only resultant values higher than the magnitude of measurement errors of negative controls were retained. Bulk atmospheric gas oxidation rate for each sample was calculated with respect to mean atmospheric mixing ratio of corresponding trace gases (H_2_: 0.53 ppmv; CO: 0.09 ppmv; CH_4_: 1.9 ppmv) ^73,145,146^. Soil cell abundance was estimated using 16S rRNA gene copy number from qPCR corrected with a reported average number of 16S rRNA gene copy per genome (i.e. 4.2) ^147^. Cell specific gas oxidation rates were then inferred by dividing estimated soil cell abundance and the proportion of corresponding gas oxidizers from metagenomic data. To identify factors potentially influencing gas oxidation rates, a two-tailed all-vs-all Spearman correlation matrix was generated that encompassed gas oxidation rates, gas oxidation gene abundances, and soil physicochemical variables for each of the 16 samples.

## Footnotes

### Etymological information

*Candidatus* Edaphomicrobium (E.da.pho.mi.cro’bi.um. Gr. neut. n. *edaphos*, soil; N.L. neut. n. *microbium*, a microbe; N. L. neut. n. *Edaphomicrobium*, a soil microbium)

*Candidatus* Edaphomicrobium janssenii (jans.sen’i.i. N.L. gen. n. *janssenii*, of Janssen, named after Peter H. Janssen, for his pioneering isolation-based studies that first described this lineage ^148^)

*Candidatus* Edaphomicrobiaceae (former candidate Chloroflexota family CSP1-4) (E.da.pho.mi.cro.bi.a.ce’ae. N.L. neut. n. *Edaphomicrobium* a (Candidatus) bacterial genus; suff. -*aceae* ending to denote a family; N.L. fem. pl. n. *Edaphomicrobiaceae*, family of the genus *Edaphomicrobium*)

*Candidatus* Edaphomicrobiales (former candidate Chloroflexota order CSP1-4) (E.da.pho.mi.cro.bi.a’les. N.L. neut. n. *Edaphomicrobium* a (Candidatus) bacterial genus; suff. -*ales* ending to denote an order; N.L. fem. pl. n. *Edaphomicrobiales*, order of the family *Edaphomicrobiaceae*)

*Candidatus* Edaphomicrobia (former candidate Chloroflexota class Ellin6529) (E.da.pho.mi.cro’bi.a. N.L. neut. n. *Edaphomicrobium* a (Candidatus) bacterial genus; -*ia* ending to denote a class; N.L. neut. pl. n. *Edaphomicrobia*, class of the order *Edaphomicrobiales*)

*Candidatus* Aridivita (A.ri.di.vi’ta. L. masc. adj. *aridus*, dry; L. fem. n. *vita*, life; N.L. fem. n. *Aridivita*, a dry life)

*Candidatus* Aridivita willemsiae (wil.lems’i.ae. N.L. gen. n. *willemsiae*, of Willems, named after Anne Willems, for her contributions to Antarctic microbiology using isolation-based approaches)

*Candidatus* Aridivitaceae (A.ri.di.vi.ta.ce’ae. N.L. neut. n. *Aridivita* a (Candidatus) bacterial genus; suff. -aceae ending to denote a family; N.L. fem. pl. n. *Aridivitaceae*, family of the genus *Aridivita*)

*Candidatus* Aridivitales (A.ri.di.vi.ta’les. N.L. neut. n. *Aridivita* a (Candidatus) bacterial genus; -ales ending to denote an order; N.L. fem. pl. n. *Aridivitales*, order of the family *Aridivitaceae*)

*Candidatus* Aridivitia (former candidate Actinobacteriota class UBA4738) (A.ri.di.vi’ti.a. N.L. neut. n. *Aridivita* a (Candidatus) bacterial genus; -ia ending to denote a class; N.L. neut. pl. n. *Aridivitia*, class of the order *Aridivitales*)

## Data availability statement

All amplicon sequencing data, raw metagenomes, and metagenome-assembled genomes were deposited to the NCBI Sequence Read Archive under BioProject accession PRJNA630822.

## Acknowledgements

This study was supported by an ARC DECRA Fellowship (DE170100310; awarded to C.G.), an Australian Antarctic Division grant (4592; awarded to C.G. and S.L.C.), a South African National Antarctic Program grant (110730: awarded to D.A.C), an NHMRC EL2 Fellowship (APP1178715; salary for C.G.), an Australian Government Research Training Stipend Scholarship and a Monash International Tuition Scholarship (awarded to P.M.L. and S.K.B.), and a National Research Foundation SANAP postdoctoral grant (awarded to M.O.). Logistic and financial support for the field work was provided by Antarctica New Zealand and the New Zealand Antarctic Research Institute, respectively (awarded to I.D.H.). We acknowledge the PhD student research discount offered by the Environmental Analysis Laboratory (EAL), a Southern Cross University NATA (National Association of Testing Authorities) ISO17025 accredited commercial and research support facility. We thank Thanavit Jirapanjawat for technical support, Maria Chuvochina for etymological advice, and Andrew Lovering, Philipp Nauer, Eleonora Chiri, Ricardo Cavicchioli, and Belinda Ferrari for helpful discussions.

## Author contributions

D.A.C., C.G., M.O., S.L.C., and I.D.H. conceived this study. C.G. and D.A.C. supervised this study. C.G., P.M.L., D.A.C., and M.O. designed experiments. P.M.L. and G.S. performed experiments. P.M.L., C.G., R.G., G.S., M.O., and D.A.C. analyzed data. P.M.L., C.G., R.G., D.A.C., M.O., and S.L.C. wrote the manuscript with input from all authors. Different authors were specifically responsible for the original sampling campaign (D.A.C., I.D.H.), metagenomic sequencing and assembly (P.M.L., C.G.), community analysis (P.M.L., G.S., C.G.), metabolic annotation (P.M.L., C.G., M.O., D.A.C.), phylogenetic analysis (C.G., P.M.L., M.O., D.A.C., S.K.B.), genetic organization analysis (P.M.L., R.G., C.G., M.O., D.A.C.), molecular modelling (R.G., C.G.), biogeochemical analysis (P.M.L., C.G., G.S.), and physicochemical analysis (P.M.L., C.G.). K.J., S.V., M.V.G. and T.M. provided theoretical and logistical support.

The authors declare no conflict of interest.

## Notes

### Competing Interest Statement

The authors have declared no competing interest.

## References

1. Leihy, R. I. et al. Antarctica’s wilderness fails to capture continent’s biodiversity. Nature 583, 567–571 (2020).

2. Cary, S. C., McDonald, I. R., Barrett, J. E. & Cowan, D. A. On the rocks: the microbiology of Antarctic Dry Valley soils. Nat. Rev. Microbiol. 8, 129–138 (2010).

3. Convey, P. et al. The spatial structure of Antarctic biodiversity. Ecol. Monogr. 84, 203–244 (2014).

4. Cavicchioli, R. On the concept of a psychrophile. ISME J. 10, 793–795 (2016).

5. Chown, S. L. et al. The changing form of Antarctic biodiversity. Nature 522, 431–438 (2015).

6. Cowan, D. A., Russell, N. J., Mamais, A. & Sheppard, D. M. Antarctic Dry Valley mineral soils contain unexpectedly high levels of microbial biomass. Extremophiles 6, 431–436 (2002).

7. Smith, J. J., Tow, L. A., Stafford, W., Cary, C. & Cowan, D. A. Bacterial diversity in three different Antarctic Cold Desert mineral soils. Microb. Ecol. 51, 413–421 (2006).

8. Lee, C. K., Barbier, B. A., Bottos, E. M., McDonald, I. R. & Cary, S. C. The Inter-Valley Soil Comparative Survey: the ecology of Dry Valley edaphic microbial communities. ISME J. 6, 1046–1057 (2012).

9. Ji, M. et al. Microbial diversity at Mitchell Peninsula, Eastern Antarctica: a potential biodiversity “hotspot”. Polar Biol. 33, 237–249 (2015).

10. Lambrechts, S., Willems, A. & Tahon, G. Uncovering the uncultivated majority in Antarctic soils: toward a synergistic approach. Front. Microbiol. 10, 242 (2019).

11. Pointing, S. B. et al. Highly specialized microbial diversity in hyper-arid polar desert. Proc. Natl. Acad. Sci. U. S. A. 107, 1254–1254 (2009).

12. Ji, M. et al. Atmospheric trace gases support primary production in Antarctic desert surface soil. Nature 552, 400–403 (2017).

13. Chan, Y., Van Nostrand, J. D., Zhou, J., Pointing, S. B. & Farrell, R. L. Functional ecology of an Antarctic Dry Valley. Proc. Natl. Acad. Sci. U. S. A. 110, 8990–8995 (2013).

14. Peeters, K., Ertz, D. & Willems, A. Culturable bacterial diversity at the Princess Elisabeth station (Utsteinen, Sør Rondane Mountains, East Antarctica) harbours many new taxa. Syst. Appl. Microbiol. 34, 360–367 (2011).

15. Pudasaini, S. et al. Microbial diversity of browning Peninsula, Eastern Antarctica revealed using molecular and cultivation methods. Front. Microbiol. 8, 591 (2017).

16. Tahon, G. & Willems, A. Isolation and characterization of aerobic anoxygenic phototrophs from exposed soils from the Sør Rondane Mountains, East Antarctica. Syst. Appl. Microbiol. 40, 357–369 (2017).

17. Tahon, G., Tytgat, B., Lebbe, L., Carlier, A. & Willems, A. *Abditibacterium utsteinense* sp. nov., the first cultivated member of candidate phylum FBP, isolated from ice-free Antarctic soil samples. Syst. Appl. Microbiol. 41, 279–290 (2018).

18. Leung, P. M. et al. Energetic basis of microbial growth and persistence in desert ecosystems. mSystems 5, e00495–19 (2020).

19. Jones, S. E. & Lennon, J. T. Dormancy contributes to the maintenance of microbial diversity. Proc. Natl. Acad. Sci. U. S. A. 107, 5881–5886 (2010).

20. Lennon, J. T. & Jones, S. E. Microbial seed banks: the ecological and evolutionary implications of dormancy. Nat. Rev. Microbiol. 9, 119–130 (2011).

21. Rittershaus, E. S. C., Baek, S. H. & Sassetti, C. M. The normalcy of dormancy: common themes in microbial quiescence. Cell Host and Microbe vol. 13 643–651 (2013).

22. Hoehler, T. M. & Jorgensen, B. B. Microbial life under extreme energy limitation. Nat. Rev. Microbiol. 11, 83–94 (2013).

23. Cockell, C. S. & Stokes, M. D. Ecology: Widespread colonization by polar hypoliths. Nature 431, 414 (2004).

24. Bay, S., Ferrari, B. & Greening, C. Life without water: How do bacteria generate biomass in desert ecosystems? Microbiol. Aust. 39, (2018).

25. Niederberger, T. D. et al. Carbon-fixation rates and associated microbial communities residing in arid and ephemerally Wet Antarctic Dry Valley soils. Front. Microbiol. 6, 1347 (2015).

26. Tahon, G., Tytgat, B., Stragier, P. & Willems, A. Analysis of *cbbL, nifH*, and *pufLM* in Soils from the Sør Rondane Mountains, Antarctica, Reveals a Large Diversity of Autotrophic and Phototrophic Bacteria. Microb. Ecol. 71, 131–149 (2016).

27. Tahon, G., Tytgat, B. & Willems, A. Diversity of key genes for carbon and nitrogen fixation in soils from the Sør Rondane Mountains, East Antarctica. Polar Biol. 41, 2181–2198 (2018).

28. Magalhães, C., Machado, A., Frank-Fahle, B., Lee, C. K. & Cary, C. S. The ecological dichotomy of ammonia-oxidizing archaea and bacteria in the hyper-arid soils of the Antarctic Dry Valleys. Front. Microbiol. 5, 515 (2014).

29. Tahon, G., Tytgat, B. & Willems, A. Diversity of phototrophic genes suggests multiple bacteria may be able to exploit sunlight in exposed soils from the Sør Rondane Mountains, East Antarctica. Front. Microbiol. 7, 2026 (2016).

30. Edwards, C. R. et al. Draft genome sequence of uncultured upland soil cluster *Gammaproteobacteria* gives molecular insights into high-affinity methanotrophy. Genome Announc. 5, e00047–17 (2017).

31. Doran, P. T. et al. Antarctic climate cooling and terrestrial ecosystem response. Nature 415, 517–520 (2002).

32. Fountain, A. G., Nylen, T. H., Monaghan, A., Basagic, H. J. & Bromwich, D. Snow in the McMurdo dry valleys, Antarctica. Int. J. Climatol. A J. R. Meteorol. Soc. 30, 633–642 (2010).

33. Van Goethem, M. W. et al. A reservoir of ‘historical’ antibiotic resistance genes in remote pristine Antarctic soils. Microbiome 6, 40 (2018).

34. Elberling, B. et al. Distribution and dynamics of soil organic matter in an Antarctic dry valley. Soil Biol. Biochem. 38, 3095–3106 (2006).

35. Adriaenssens, E. M. et al. Environmental drivers of viral community composition in Antarctic soils identified by viromics. Microbiome 5, 83 (2017).

36. Bay, S. K. et al. Soil bacterial communities exhibit strong biogeographic patterns at fine taxonomic resolution. mSystems 5, e00540–20 (2020).

37. Janssen, P. H. Identifying the dominant soil bacterial taxa in libraries of 16S rRNA and 16S rRNA genes. Appl. Environ. Microbiol. 72, 1719–1728 (2006).

38. Delgado-Baquerizo, M. et al. A global atlas of the dominant bacteria found in soil. Science 359, 320–325 (2018).

39. Rotem, O., Pasternak, Z. & Jurkevitch, E. The genus Bdellovibrio and like organisms. in The Prokaryotes: Deltaproteobacteria and Epsilonproteobacteria vol. 9783642390 3–17 (2014).

40. Collingro, A., Köstlbacher, S. & Horn, M. Chlamydiae in the Environment. Trends Microbiol. 10.1016/j.tim.2020.05.020 (2020) doi: 10.1016/j.tim.2020.05.020.

41. Yeoh, Y. K., Sekiguchi, Y., Parks, D. H. & Hugenholtz, P. Comparative genomics of candidate phylum TM6 suggests that parasitism is widespread and ancestral in this lineage. Mol. Biol. Evol. 33, 915–927 (2016).

42. Brown, C. T. et al. Unusual biology across a group comprising more than 15% of domain *Bacteria*. Nature 523, 208–211 (2015).

43. Castelle, C. J. & Banfield, J. F. Major new microbial groups expand diversity and alter our understanding of the tree of life. Cell 172, 1181–1197 (2018).

44. Bowers, R. M. et al. Minimum information about a single amplified genome (MISAG) and a metagenome-assembled genome (MIMAG) of bacteria and archaea. Nat. Biotechnol. 35, 725–731 (2017).

45. Constant, P., Chowdhury, S. P., Pratscher, J. & Conrad, R. Streptomycetes contributing to atmospheric molecular hydrogen soil uptake are widespread and encode a putative high-affinity [NiFe]-hydrogenase. Environ. Microbiol. 12, 821–829 (2010).

46. Greening, C., Berney, M., Hards, K., Cook, G. M. & Conrad, R. A soil actinobacterium scavenges atmospheric H2 using two membrane-associated, oxygen-dependent [NiFe] hydrogenases. Proc. Natl. Acad. Sci. U. S. A. 111, 4257–4261 (2014).

47. Greening, C. et al. Persistence of the dominant soil phylum Acidobacteria by trace gas scavenging. Proc. Natl. Acad. Sci. U. S. A. 112, 10497–10502 (2015).

48. Søndergaard, D., Pedersen, C. N. S. & Greening, C. HydDB: a web tool for hydrogenase classification and analysis. Sci. Rep. 6, 34212 (2016).

49. Constant, P., Chowdhury, S. P., Hesse, L., Pratscher, J. & Conrad, R. Genome data mining and soil survey for the novel Group 5 [NiFe]-hydrogenase to explore the diversity and ecological importance of presumptive high-affinity H2-oxidizing bacteria. Appl. Environ. Microbiol. 77, 6027–6035 (2011).

50. King, G. M. Molecular and culture-based analyses of aerobic carbon monoxide oxidizer diversity. Appl. Environ. Microbiol. 69, 7257–7265 (2003).

51. Islam, Z. F. et al. Two Chloroflexi classes independently evolved the ability to persist on atmospheric hydrogen and carbon monoxide. ISME J. 13, 1801–1813 (2019).

52. Cordero, P. R. F. et al. Atmospheric carbon monoxide oxidation is a widespread mechanism supporting microbial survival. ISME J. 13, 2868–2881 (2019).

53. King, G. M. & Weber, C. F. Distribution, diversity and ecology of aerobic CO-oxidizing bacteria. Nat. Rev. Microbiol. 5, 107–118 (2007).

54. Tabita, F. R. et al. Function, structure, and evolution of the RubisCO-like proteins and their RubisCO homologs. Microbiol. Mol. Biol. Rev. 71, 576–599 (2007).

55. Park, S. W. et al. Presence of duplicate genes encoding a phylogenetically new subgroup of form I ribulose 1,5-bisphosphate carboxylase/oxygenase in *Mycobacterium* sp. strain JC1 DSM 3803. Res. Microbiol. 160, 159–165 (2009).

56. Grostern, A. & Alvarez-Cohen, L. RubisCO-based CO2 fixation and C1 metabolism in the actinobacterium *Pseudonocardia dioxanivorans* CB1190. Environ. Microbiol. 15, 3040–3053 (2013).

57. Greening, C., Villas-Bôas, S. G., Robson, J. R., Berney, M. & Cook, G. M. The growth and survival of *Mycobacterium smegmatis* is enhanced by co-metabolism of atmospheric H2. PLoS One 9, e103034 (2014).

58. Liot, Q. & Constant, P. Breathing air to save energy – new insights into the ecophysiological role of high-affinity [NiFe]-hydrogenase in Streptomyces avermitilis. Microbiologyopen 5, 47–59 (2016).

59. Tveit, A. T. et al. Widespread soil bacterium that oxidizes atmospheric methane. Proc. Natl. Acad. Sci. U. S. A. 116, 8515–8524 (2019).

60. Islam, Z. F. et al. A widely distributed hydrogenase oxidises atmospheric H2 during bacterial growth. ISME J. 10.1038/s41396-020-0713–4 (2020) doi: 10.1101/2020.04.14.040717.

61. Schmitz, R. A. et al. The thermoacidophilic methanotroph *Methylacidiphilum fumariolicum* SolV oxidizes subatmospheric H2 with a high-affinity, membrane-associated [NiFe] hydrogenase. ISME J. 14, 1223–1232 (2020).

62. Cordero, P. R. F. et al. Two uptake hydrogenases differentially interact with the aerobic respiratory chain during mycobacterial growth and persistence. J. Biol. Chem. 294, 18980–18991 (2019).

63. Lopes, A. R., Manaia, C. M. & Nunes, O. C. Bacterial community variations in an alfalfa-rice rotation system revealed by 16S rRNA gene 454-pyrosequencing. FEMS Microbiol. Ecol. 87, 650–663 (2014).

64. Frindte, K., Pape, R., Werner, K., Löffler, J. & Knief, C. Temperature and soil moisture control microbial community composition in an arctic–alpine ecosystem along elevational and micro-topographic gradients. ISME J. 13, 2031–2043 (2019).

65. Khilyas, I. V. et al. Microbial diversity and mineral composition of weathered serpentine rock of the Khalilovsky massif. PLoS One 14, e0225929 (2019).

66. Johnston, E. R. et al. Responses of tundra soil microbial communities to half a decade of experimental warming at two critical depths. Proc. Natl. Acad. Sci. 116, 15096–15105 (2019).

67. Whitman, W. B. Modest proposals to expand the type material for naming of prokaryotes. Int. J. Syst. Evol. Microbiol. 66, 2108–2112 (2016).

68. Murray, A. E. et al. Roadmap for naming uncultivated Archaea and Bacteria. Nat. Microbiol. 5, 987–994 (2020).

69. Hirsch, P. et al. *Hymenobacter roseosalivarius* gen. nov., sp. nov. from continental Antarctic soils and sandstone: bacteria of the *Cytophaga*/*Flavobacterium*/*Bacteroides* line of phylogenetic descent. Syst. Appl. Microbiol. 21, 374–383 (1998).

70. Schäfer, C., Friedrich, B. & Lenz, O. Novel, oxygen-insensitive group 5 [NiFe]-hydrogenase in Ralstonia eutropha. Appl. Environ. Microbiol. 79, 5137–45 (2013).

71. Schäfer, C. et al. Structure of an actinobacterial-type [NiFe]-hydrogenase reveals insight into O2-tolerant H2 oxidation. Structure 24, 285–292 (2016).

72. Ehhalt, D. H. & Rohrer, F. The tropospheric cycle of H2: a critical review. Tellus B 61, 500–535 (2009).

73. Novelli, P. C., Masarie, K. A. & Lang, P. M. Distributions and recent changes of carbon monoxide in the lower troposphere. J. Geophys. Res. Atmos. 103, 19015–19033 (1998).

74. Conrad, R. Soil microorganisms as controllers of atmospheric trace gases (H2, CO, CH4, OCS, N2O, and NO). Microbiol. Mol. Biol. Rev. 60, 609–640 (1996).

75. Greening, C., Grinter, R. & Chiri, E. Uncovering the metabolic strategies of the dormant microbial majority: towards integrative approaches. mSystems 4, e00107–19 (2019).

76. Tijhuis, L., Van Loosdrecht, M. C. & Heijnen, J. J. A thermodynamically based correlation for maintenance gibbs energy requirements in aerobic and anaerobic chemotrophic growth. Biotechnol. Bioeng. 42, 509–519 (1993).

77. Price, P. B. & Sowers, T. Temperature dependence of metabolic rates for microbial growth, maintenance, and survival. Proc. Natl. Acad. Sci. U. S. A. 101, 4631–4636 (2004).

78. LaRowe, D. E. & Amend, J. P. Power limits for microbial life. Front. Microbiol. 6, 718 (2015).

79. Sundararaj, S. et al. The CyberCell Database (CCDB): A comprehensive, self-updating, relational database to coordinate and facilitate in silico modeling of Escherichia coli. Nucleic Acids Res. 32, D293–D295 (2004).

80. Koch, A. L. What size should a bacterium be? A question of scale. Annu. Rev. Microbiol. 50, 317–348 (1996).

81. Finkel, O. M., Béja, O. & Belkin, S. Global abundance of microbial rhodopsins. ISME J. 7, 448–451 (2013).

82. Pinhassi, J., DeLong, E. F., Béjà, O., González, J. M. & Pedrós-Alió, C. Marine bacterial and archaeal ion-pumping rhodopsins: genetic Diversity, physiology, and ecology. Microbiol. Mol. Biol. Rev. 80, 929–954 (2016).

83. Gomez-Consarnau, L. et al. Proteorhodopsin Phototrophy Promotes Survival of Marine Bacteria during Starvation. Plos Biol. 8, (2010).

84. Steindler, L., Schwalbach, M. S., Smith, D. P., Chan, F. & Giovannoni, S. J. Energy starved candidatus *Pelagibacter ubique* substitutes light-mediated ATP production for endogenous carbon respiration. PLoS One 6, e19725 (2011).

85. Panwar, P. et al. Influence of the polar light cycle on seasonal dynamics of an Antarctic lake microbial community. Microbiome 10.1186/s40168-020-00889–8 (2020).

86. Guerrero, L. D., Vikram, S., Makhalanyane, T. P. & Cowan, D. A. Evidence of microbial rhodopsins in Antarctic Dry Valley edaphic systems. Environ. Microbiol. 19, 3755–3767 (2017).

87. Oesterhelt, D. & Stoeckenius, W. Rhodopsin-like protein from the purple membrane of *Halobacterium halobium*. Nat. new Biol. 233, 149–152 (1971).

88. Harris, A. et al. A new group of eubacterial light-driven retinal-binding proton pumps with an unusual cytoplasmic proton donor. Biochim. Biophys. Acta (BBA)-Bioenergetics 1847, 1518–1529 (2015).

89. Deeg, C. M. et al. *Chromulinavorax destructans*, a pathogen of microzooplankton that provides a window into the enigmatic candidate phylum Dependentiae. PLoS Pathog. 15, e1007801 (2019).

90. Beet, C. R. et al. Genetic diversity among populations of Antarctic springtails (Collembola) within the Mackay Glacier ecotone. Genome 59, 762–770 (2016).

91. Lambert, C. et al. Ankyrin-mediated self-protection during cell invasion by the bacterial predator *Bdellovibrio bacteriovorus*. Nat. Commun. 6, 1–10 (2015).

92. Pasternak, Z. et al. By their genes ye shall know them: genomic signatures of predatory bacteria. ISME J. 7, 756–769 (2013).

93. Hamm, J. N. et al. Unexpected host dependency of Antarctic Nanohaloarchaeota. Proc. Natl. Acad. Sci. 116, 14661–14670 (2019).

94. Lagkouvardos, I. et al. Integrating metagenomic and amplicon databases to resolve the phylogenetic and ecological diversity of the Chlamydiae. ISME J. 8, 115–125 (2014).

95. Jaffe, A. L., Castelle, C. J., Carnevali, P. B. M., Gribaldo, S. & Banfield, J. F. The rise of diversity in metabolic platforms across the Candidate Phyla Radiation. BMC Biol. 18, 1–15 (2020).

96. Beam, J. P. et al. Ancestral absence of electron transport chains in Patescibacteria and DPANN. bioRxiv 2020.04.07.029462 (2020) doi: 10.1101/2020.04.07.029462.

97. Cowan, D. A. & Makhalanyane, T. P. Energy from thin air. Nature 552, 336–337 (2017).

98. Lee, J. R. et al. Climate change drives expansion of Antarctic ice-free habitat. Nature 547, 49 (2017).

99. Rintoul, S. R. et al. Choosing the future of Antarctica. Nature 558, 233–241 (2018).

100. Cavicchioli, R. et al. Scientists’ warning to humanity: microorganisms and climate change. Nat. Rev. Microbiol. 17, 569–586 (2019).

101. Kennicutt, M. C. 2nd et al. Six priorities for Antarctic science. Nature 512, 23–25 (2014).

102. Heldmann, J. L. et al. The high elevation Dry Valleys in Antarctica as analog sites for subsurface ice on Mars. Planet. Space Sci. 85, 53–58 (2013).

103. Nurk, S., Meleshko, D., Korobeynikov, A. & Pevzner, P. A. metaSPAdes: a new versatile metagenomic assembler. Genome Res. 27, 824–834 (2017).

104. Li, D. H. et al. MEGAHIT v1.0: A fast and scalable metagenome assembler driven by advanced methodologies and community practices. Methods 102, 3–11 (2016).

105. Langmead, B. & Salzberg, S. L. Fast gapped-read alignment with Bowtie 2. Nat. Methods 9, 357 (2012).

106. Alneberg, J. et al. Binning metagenomic contigs by coverage and composition. Nat. Methods 11, 1144 (2014).

107. Wu, Y.-W., Simmons, B. A. & Singer, S. W. MaxBin 2.0: an automated binning algorithm to recover genomes from multiple metagenomic datasets. Bioinformatics 32, 605–607 (2015).

108. Kang, D. et al. MetaBAT 2: an adaptive binning algorithm for robust and efficient genome reconstruction from metagenome assemblies. PeerJ 7, e7359 (2019).

109. Sieber, C. M. K. et al. Recovery of genomes from metagenomes via a dereplication, aggregation and scoring strategy. Nat. Microbiol. 1 (2018).

110. Parks, D. H. et al. Recovery of nearly 8,000 metagenome-assembled genomes substantially expands the tree of life. Nat. Microbiol. 2, 1533 (2017).

111. Olm, M. R., Brown, C. T., Brooks, B. & Banfield, J. F. dRep: a tool for fast and accurate genomic comparisons that enables improved genome recovery from metagenomes through de-replication. ISME J. 11, 2864 (2017).

112. Parks, D. H., Imelfort, M., Skennerton, C. T., Hugenholtz, P. & Tyson, G. W. CheckM: assessing the quality of microbial genomes recovered from isolates, single cells, and metagenomes. Genome Res. 25, 1043–1055 (2015).

113. Chaumeil, P.-A., Mussig, A. J., Hugenholtz, P. & Parks, D. H. GTDB-Tk: a toolkit to classify genomes with the Genome Taxonomy Database. Bioinformatics 36, 1925–1927 (2020).

114. Parks, D. H. et al. A standardized bacterial taxonomy based on genome phylogeny substantially revises the tree of life. Nat. Biotechnol. 36, 996–1004 (2018).

115. Hyatt, D. et al. Prodigal: prokaryotic gene recognition and translation initiation site identification. BMC Bioinformatics 11, 119 (2010).

116. Boyd, J. A., Woodcroft, B. J. & Tyson, G. W. GraftM: a tool for scalable, phylogenetically informed classification of genes within metagenomes. Nucleic Acids Res. 46, e59–e59 (2018).

117. Chen, Y.-J. et al. Metabolic flexibility allows generalist bacteria to become dominant in a frequently disturbed ecosystem. bioRxiv 2020.02.12.945220 (2020).

118. Parada, A. E., Needham, D. M. & Fuhrman, J. A. Every base matters: assessing small subunit rRNA primers for marine microbiomes with mock communities, time series and global field samples. Environ. Microbiol. 18, 1403–1414 (2016).

119. Apprill, A., McNally, S., Parsons, R. & Weber, L. Minor revision to V4 region SSU rRNA 806R gene primer greatly increases detection of SAR11 bacterioplankton. Aquat. Microb. Ecol. 75, 129–137 (2015).

120. Bolyen, E. et al. Reproducible, interactive, scalable and extensible microbiome data science using QIIME 2. Nat. Biotechnol. 37, 852–857 (2019).

121. Martin, M. Cutadapt removes adapter sequences from high-throughput sequencing reads. EMBnet.journal 17, 10 (2011).

122. Rognes, T., Flouri, T., Nichols, B., Quince, C. & Mahé, F. VSEARCH: a versatile open source tool for metagenomics. PeerJ 4, 2584 (2016).

123. Amir, A. et al. Deblur rapidly resolves single-nucleotide community sequence patterns. mSystems 2, e00191–16 (2017).

124. Quast, C. et al. The SILVA ribosomal RNA gene database project: improved data processing and web-based tools. Nucleic Acids Res. 41, D590–D596 (2012).

125. McDonald, D. et al. An improved Greengenes taxonomy with explicit ranks for ecological and evolutionary analyses of bacteria and archaea. ISME J. 6, 610–618 (2012).

126. Katoh, K., Misawa, K., Kuma, K. I. & Miyata, T. MAFFT: A novel method for rapid multiple sequence alignment based on fast Fourier transform. Nucleic Acids Res. 30, 3059–3066 (2002).

127. Price, M. N., Dehal, P. S. & Arkin, A. P. FastTree 2 - Approximately maximum-likelihood trees for large alignments. PLoS One 5, (2010).

128. McMurdie, P. J. & Holmes, S. phyloseq: an R package for reproducible interactive analysis and graphics of microbiome census data. PLoS One 8, e61217 (2013).

129. Kembel, S. W. et al. Picante: R tools for integrating phylogenies and ecology. Bioinformatics 26, 1463–1464 (2010).

130. Dixon, P. VEGAN, a package of R functions for community ecology. J. Veg. Sci. 14, 927–930 (2003).

131. Baselga, A. & Orme, C. D. L. Betapart: An R package for the study of beta diversity. Methods Ecol. Evol. 3, 808–812 (2012).

132. Wickham, H. ggplot2: elegant graphics for data analysis. (Springer, 2016).

133. Buchfink, B., Xie, C. & Huson, D. H. Fast and sensitive protein alignment using DIAMOND. Nat. Methods 12, 59 (2014).

134. Greening, C. et al. Diverse hydrogen production and consumption pathways influence methane production in ruminants. ISME J. 13, 2617–2632 (2019).

135. Pruitt, K. D., Tatusova, T. & Maglott, D. R. NCBI reference sequences (RefSeq): a curated non-redundant sequence database of genomes, transcripts and proteins. Nucleic Acids Res. 35, D61–D65 (2007).

136. Eddy, S. R. Accelerated profile HMM searches. PLoS Comput. Biol. 7, e1002195 (2011).

137. Anantharaman, K. et al. Thousands of microbial genomes shed light on interconnected biogeochemical processes in an aquifer system. Nat. Commun. 7, 13219 (2016).

138. Darling, A. E. et al. PhyloSift: phylogenetic analysis of genomes and metagenomes. PeerJ 2, e243 (2014).

139. Larkin, M. A. et al. Clustal W and Clustal X version 2.0. Bioinformatics 23, 2947–2948 (2007).

140. Kumar, S., Stecher, G., Li, M., Knyaz, C. & Tamura, K. MEGA X: molecular evolutionary genetics analysis across computing platforms. Mol. Biol. Evol. 35, 1547–1549 (2018).

141. Finn, R. D. et al. Pfam: The protein families database. Nucleic Acids Research vol. 42 D222–D230 (2014).

142. Madeira, F. et al. The EMBL-EBI search and sequence analysis tools APIs in 2019. Nucleic Acids Res. 47, W636–W641 (2019).

143. Kelley, L. A., Mezulis, S., Yates, C. M., Wass, M. N. & Sternberg, M. J. E. The Phyre2 web portal for protein modeling, prediction and analysis. Nat. Protoc. 10, 845–858 (2015).

144. Krogh, A., Larsson, B., Von Heijne, G. & Sonnhammer, E. L. L. Predicting transmembrane protein topology with a hidden Markov model: Application to complete genomes. J. Mol. Biol. 305, 567–580 (2001).

145. Novelli, P. C. et al. Molecular hydrogen in the troposphere: global distribution and budget. J. Geophys. Res. Atmos. 104, 30427–30444 (1999).

146. Stocker, T. F. et al. Climate change 2013 the physical science basis: Working Group I contribution to the fifth assessment report of the intergovernmental panel on climate change. Climate Change 2013 the Physical Science Basis: Working Group I Contribution to the Fifth Assessment Report of the Intergovernmental Panel on Climate Change vol. 9781107057 (2013).

147. Větrovský, T. & Baldrian, P. The variability of the 16S rRNA gene in bacterial genomes and its consequences for bacterial community analyses. PLoS One 8, e57923 (2013).

148. Davis, K. E. R., Sangwan, P. & Janssen, P. H. Acidobacteria, Rubrobacteridae and Chloroflexi are abundant among very slow-growing and mini-colony-forming soil bacteria. Environ. Microbiol. 13, 798–805 (2011).

