## Supplementary information for "A genome compendium reveals diverse metabolic adaptations of Antarctic soil microorganisms"

**Supplementary Note 1.** Genomic analysis and structural modelling suggest that the group 1l [NiFe]-hydrogenase is a *bona fide* H_2_-oxidizing enzyme with multiple unique features. Analysis of neighboring genes of catalytic subunit of this enzyme (HylL) showed the presence of canonical hydrogenase accessory genes including the hydrogenase small subunit (HylS), hydrogenase maturation protease (HupF), carbamoyltransferase (HypF), and a Rieske-like iron-sulfur protein (HylE) homologous to HhyE (**Figure 3b**, **Figure S5, Table S7**), supporting that these hydrogenase sequences are *bona fide*. We generated a structural model of the HylL and HylS subunits from *Hymenobacter roseosalivarius*, which was previously isolated from soils of Antarctic Dry Valleys. This model revealed that the amino acid sequence of HylL and HylS is consistent with the formation of a functional heterotetramer, analogous to that observed in group 1h [NiFe] hydrogenases (**Figure 3b**). Additionally, the C-terminus of the HylS subunit possesses a C-terminal ~100 amino acid extension, relative to group 1h hydrogenases. The residues required for [FeS] cluster and [NiFe] centre coordination by HylS and HylL respectively are conserved, validating the catalytic function of the hydrogenase (**Figure 3c, d**). The [FeS] cluster of HylS proximal to HylL is coordinated by four cysteine residues, showing divergence from group 1h [NiFe]-hydrogenases where an aspartate generally substitutes one cysteine residue. A common feature of the genetic organization of group 1l [NiFe]-hydrogenases is the presence of five small open reading frames separating the genes for the HylS and HylL subunits. While these five open reading frames share only 12-30% amino acid identity, they are all predicted to encode single-pass transmembrane proteins, which we designate HylTM1-5. Intriguingly, HylTM4 contains a C-terminal tri-cysteine motif, potentially involved in [FeS] cluster coordination. While further experimental analysis of these HylTM proteins is required, their predicted structure and membrane localization, as well as their intimate genetic association with the HylS and HylL subunits, suggests a physical interaction, possibly through the C-terminal extension of HylS (**Figure 3b**). This suggests they may constitute a novel membrane anchoring and electron relaying system for 1l [NiFe]-hydrogenases, and an alternative mechanism of metabolic integration to that of the group 1h [NiFe]-hydrogenases.

**Supplementary Note 2.** Correlational analysis suggests various physicochemical factors influence cell-specific trace gas oxidation rates. While methane monooxygenase abundance predicted methane oxidation rates (*p* = 0.013), hydrogenase and CO dehydrogenase abundance were weakly correlated with H_2_ oxidation rates (*p* = 0.139) and CO oxidation rate (*p* = 0.381) respectively (**Figure S6; Table S9**). Instead, H_2_ oxidation rate is strongly correlated with pH, salinity and associated factors, and sulfur content (*p* < 0.01), whereas CO oxidation rate appears to be moderately correlated with soil sulfur and magnesium content (*p* < 0.05). This indicates that, in addition to the presence of metabolic capacity for trace gas oxidation, biological activity could be constrained by other environmental factors. However, as soil samples used in this study spanned a large varied ecotone and were not sampled along a gradient transect, we are unable to definitively confirm these observations. Larger-scale field-based surveys and laboratory-based manipulations are required to discern the various factors driving the abundance and activities of trace gas oxidizers in Antarctic soils and other ecosystems.

**Table S1 (xlsx).** Sampling metadata, physicochemical parameters, and 16S rRNA gene copy number of soils collected from the Mackay Glacier region.

**Table S2 (xlsx).** Community composition and alpha diversity of soil bacterial and archaeal community using the metagenome universal single copy marker *rplP*.

**Table S3 (xlsx).** Sequencing details, community composition and alpha diversity of bacterial and archaeal communities based on 16S rRNA amplicon data.

**Table S4 (xlsx).** Sequencing and assembly details of the sixteen shotgun metagenomes.

**Table S5 (xlsx)**. Summary of taxonomy, quality statistics, coverage, and genetic capabilities of the 451 metagenome-assembled genomes.

**Table S6 (xlsx).** Abundance of metabolic marker genes in soil metagenome short reads.

**Table S7 (xlsx).** Annotation of neighbouring genes flanking large subunits of RuBisCO and [NiFe] hydrogenases.

**Table S8 (xlsx).** Measurements of H_2_, CH_4_, and CO concentrations in soil microcosm timecourses.

**Table S9 (xlsx).** Correlation matrix showing significance and correlation between soil physicochemical parameters, genetic determinants for trace gas oxidation and gas oxidation rates.

**Figure S1.** Rarefaction curves, community composition, alpha and beta diversity based on 16S rRNA gene amplicon sequence variants (ASVs). Rarefaction curves based on observed ASVs are shown **(a)** before rarefaction and **(b)** after rarefying to 13248 sequences for each sample. Rarefaction curves approaching asymptote indicate that sequencing depth was adequate to capture the soil microbial diversity within each sample. **(c)** Stacked bar chart showing phylum-level bacterial and archaeal community composition based on ASVs taxonomically annotated by Silva database release 138. Phyla with less than 1% abundance in the sample were grouped to “Other phyla”. **(d)** Boxplot showing alpha diversity (observed richness, estimated richness, Shannon diversity, Faith’s phylogenetic diversity) of microbial communities after rarefying 16S rRNA gene amplicon data to 13248 sequences for each sample. **(e)** Distance decay relationship of beta diversity (Bray-Curtis dissimilarity) with increasing pairwise geographic distance. An exponential regression curve and 95% confidence interval are shown.

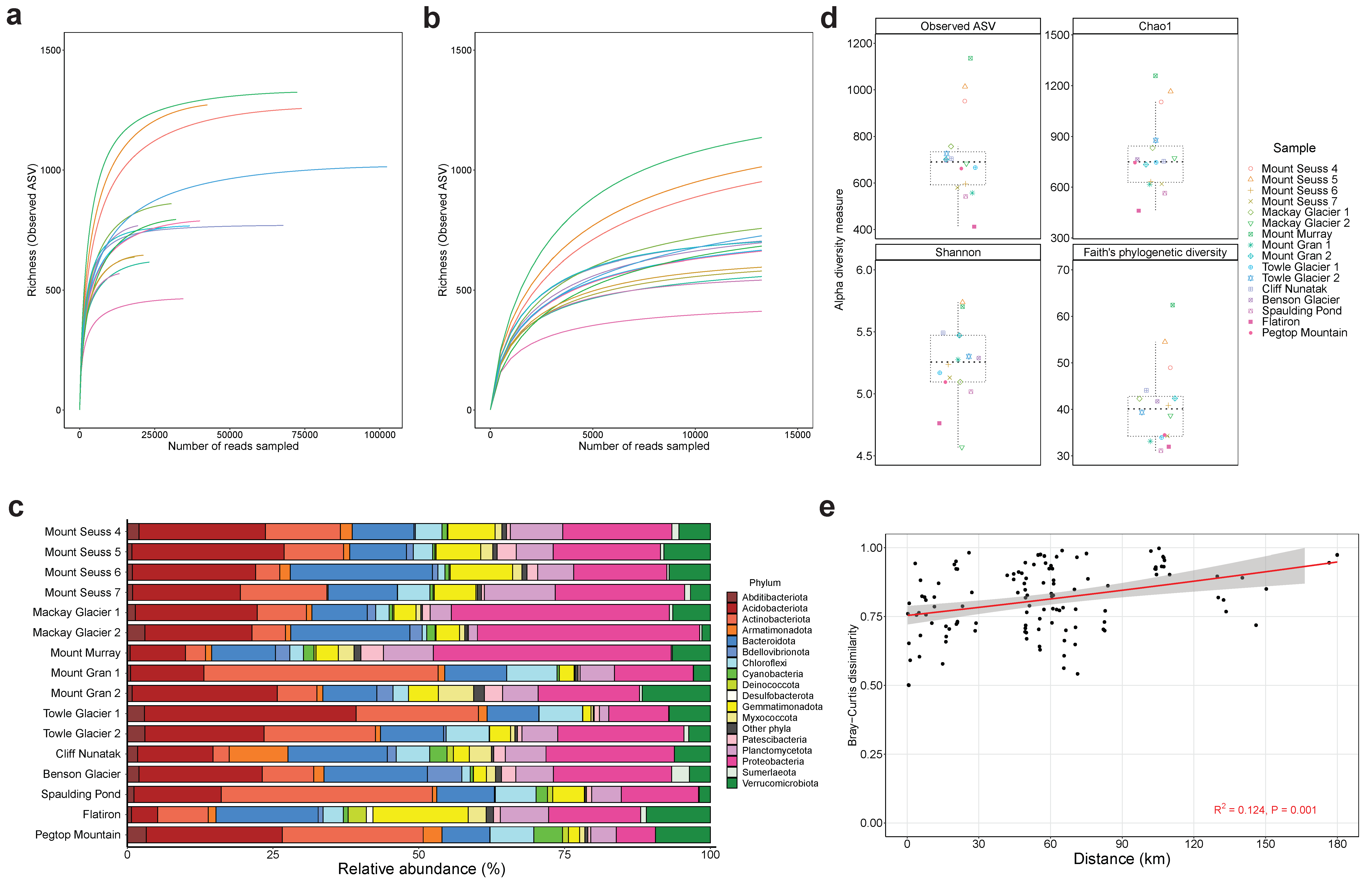

**Figure S2.** Ability to utilize inorganic compounds via reductive pathways in the microbial communities. Homology-based searches were used to identify signature genes encoding enzymes associated with (from top to bottom): methanogenesis, hydrogenogenic fermentation, sulfate reduction, nitrogen cycling pathways, iron reduction, and other processes. The left heatmap shows the percentage of total community members predicted to encode each signature metabolic gene. To infer abundance, read counts were normalized to gene length and the abundance of single-copy marker genes. The right heatmap shows the presence of these genes across the 451 metagenome-assembled genomes spanning 18 phyla. Abundance was normalized by predicted MAG completeness.

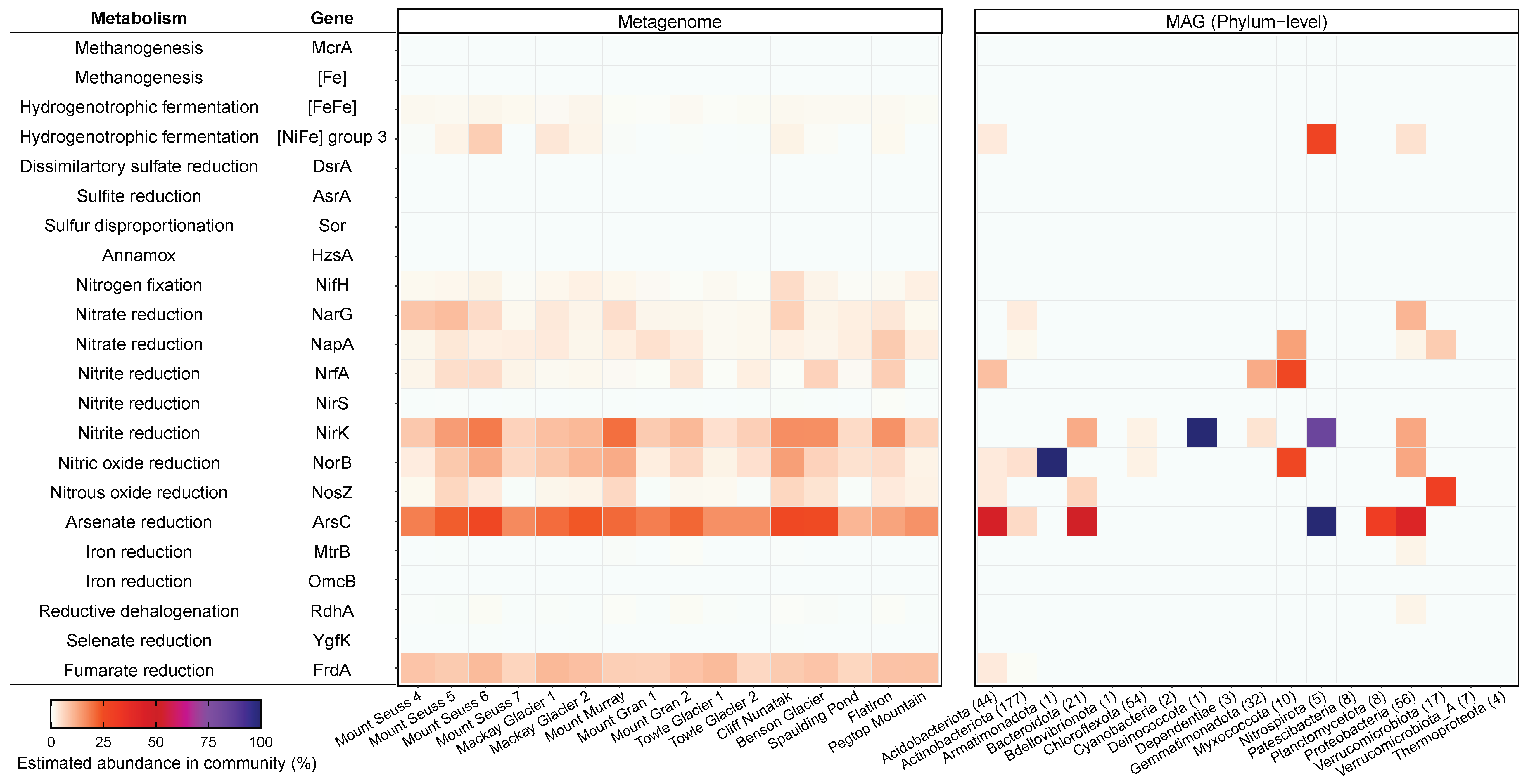

**Figure S3.** Maximum-likelihood tree of amino acid sequences of the form I [MoCu]-carbon monoxide carbon monoxide dehydrogenase catalytic subunit (CoxL), a marker for aerobic CO oxidation. The tree shows sequences from metagenome-assembled genomes (blue) from the Mackay Glacier region alongside representative reference sequences (grey). The tree was constructed using the JTT matrix-based model, used all sites, and was bootstrapped with 50 replicates and midpoint-rooted.

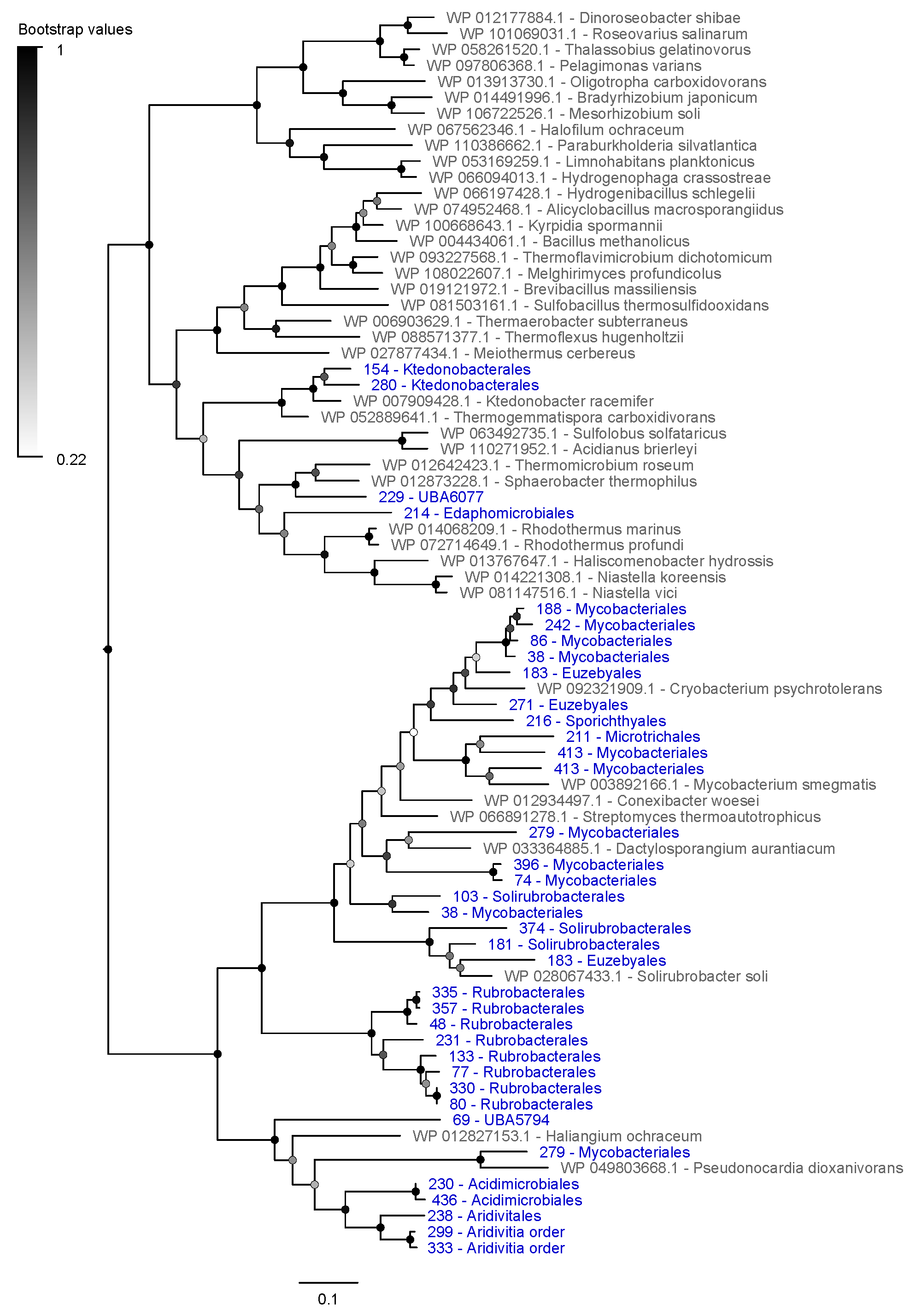

**Figure S4.** Maximum-likelihood tree of amino acid sequences of ribulose 1,5-bisphosphate carboxylase/oxygenase (RuBisCO) large subunit (RbcL), a marker for carbon fixation through the Calvin-Benson-Bassham (CBB) cycle. The tree shows sequences from metagenome-assembled genomes (blue) from the Mackay Glacier region alongside representative reference sequences (grey). The RuBisCO subtype (IA to IE, II, III) is shown next to the reference sequences. The tree was constructed using the JTT matrix-based model, used all sites, and was bootstrapped with 50 replicates and midpoint-rooted.

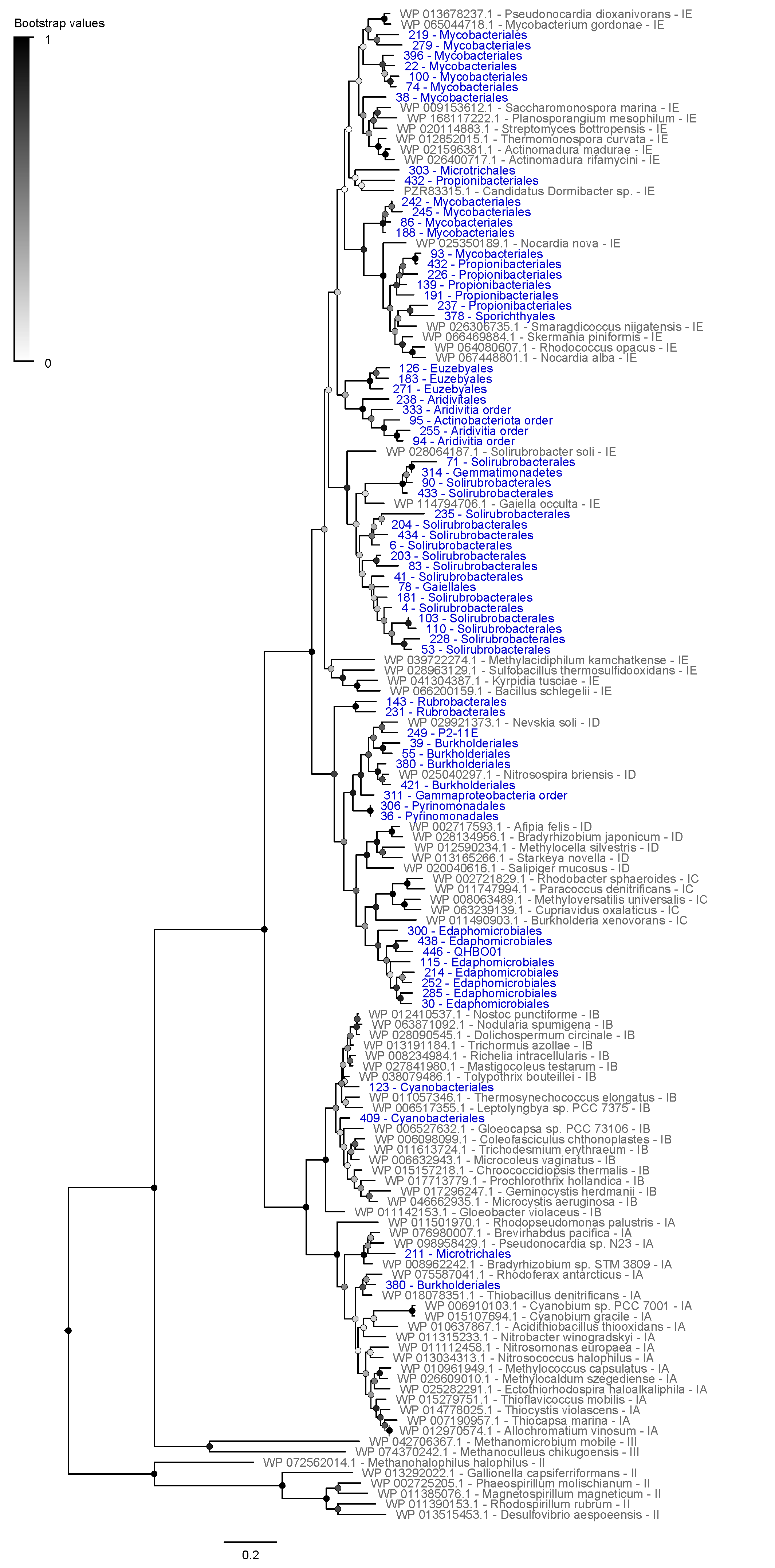

**Figure S5.** Comparison of genetic organization of (**a**) group 1h and (**b**) group 1l [NiFe] hydrogenases from metagenome-assembled genomes and representative bacterial isolate genomes. Up to 10 genes upstream and downstream of the hydrogenase large subunits were shown. Number and content in brackets denote MAG identifier and their corresponding taxonomy at the order level, respectively. Abbreviations: CbiA = CobQ/CobB/MinD/ParA nucleotide binding domain containing protein; DUF1059 = Protein of unknown function (DUF1059); HhyE = Electron-relaying Rieske-type protein of group 1h hydrogenase; HhyL = Large subunits of group 1h hydrogenase; HhyS = Small subunits of group 1h hydrogenase; HUO = Hypothetical proteins, proteins with Unknown function or Other proteins; HupD = hydrogenase maturation endopeptidase; HylE = Electron-relaying Rieske-type protein of group 1l hydrogenase; HylL = Large subunits of group 1l hydrogenase; HylS = Small subunits of group 1l hydrogenase; HylTM = Single-pass transmembrane proteins; HypABCDEF = Hydrogenase maturation factors; PadR = PadR family transcriptional regulator; PF00497 = Bacterial extracellular solute-binding proteins, family 3; PF00528 = Binding-protein-dependent transport system; SIS = Phosphoglucose isomerase; TNR = Tetratricopeptide repeat protein.

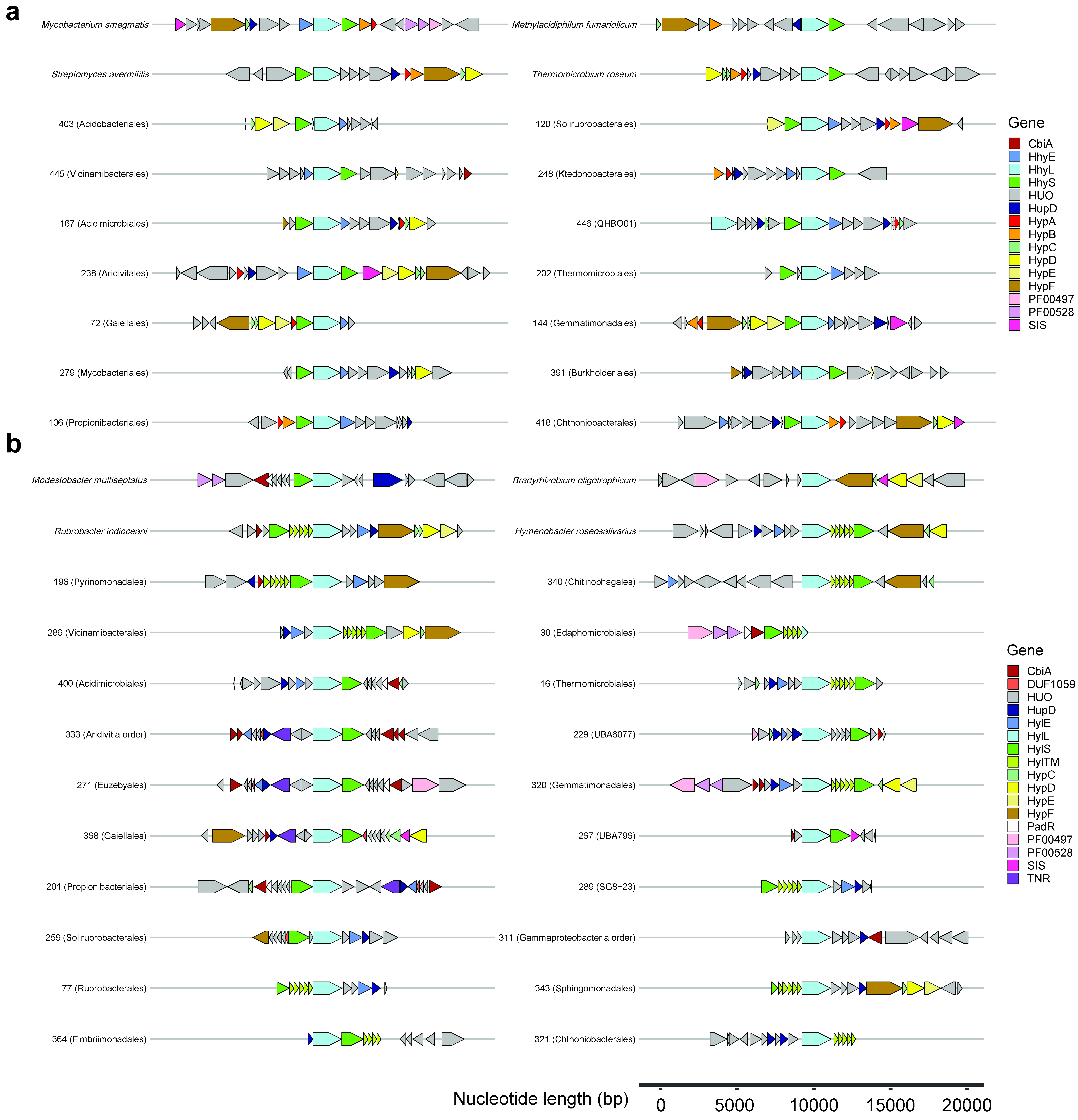

**Figure S6.** Heatmap showing correlations between soil physicochemical parameters, microbial signature markers, and biogeochemical activities. 19 representative soil physicochemical parameters are shown: elevation, moisture content, pH, electrical conductivity, carbon/nitrogen ratio, total organic carbon, nitrate, ammonium, sulfur, phosphorus, effective cation exchange capacity (ECEC), exchangeable calcium, magnesium, potassium, sodium, manganese, iron, copper, chloride. Also shown are seven microbial signature markers: 16S rRNA gene copy number and abundance of group 1h [NiFe] hydrogenase (HhyL), group 1l [NiFe] hydrogenase (HylL), all [NiFe] hydrogenases, carbon monoxide dehydrogenase (CoxL), methane monooxygenases (PmoA & MmoA), and RuBisCO (RbcL). Finally the bulk oxidation and cell-specific rates of atmospheric H_2_, CO, and CH_4_ oxidation were included. Red gradient indicates negative correlations. Blue gradient reflects more positive correlations.

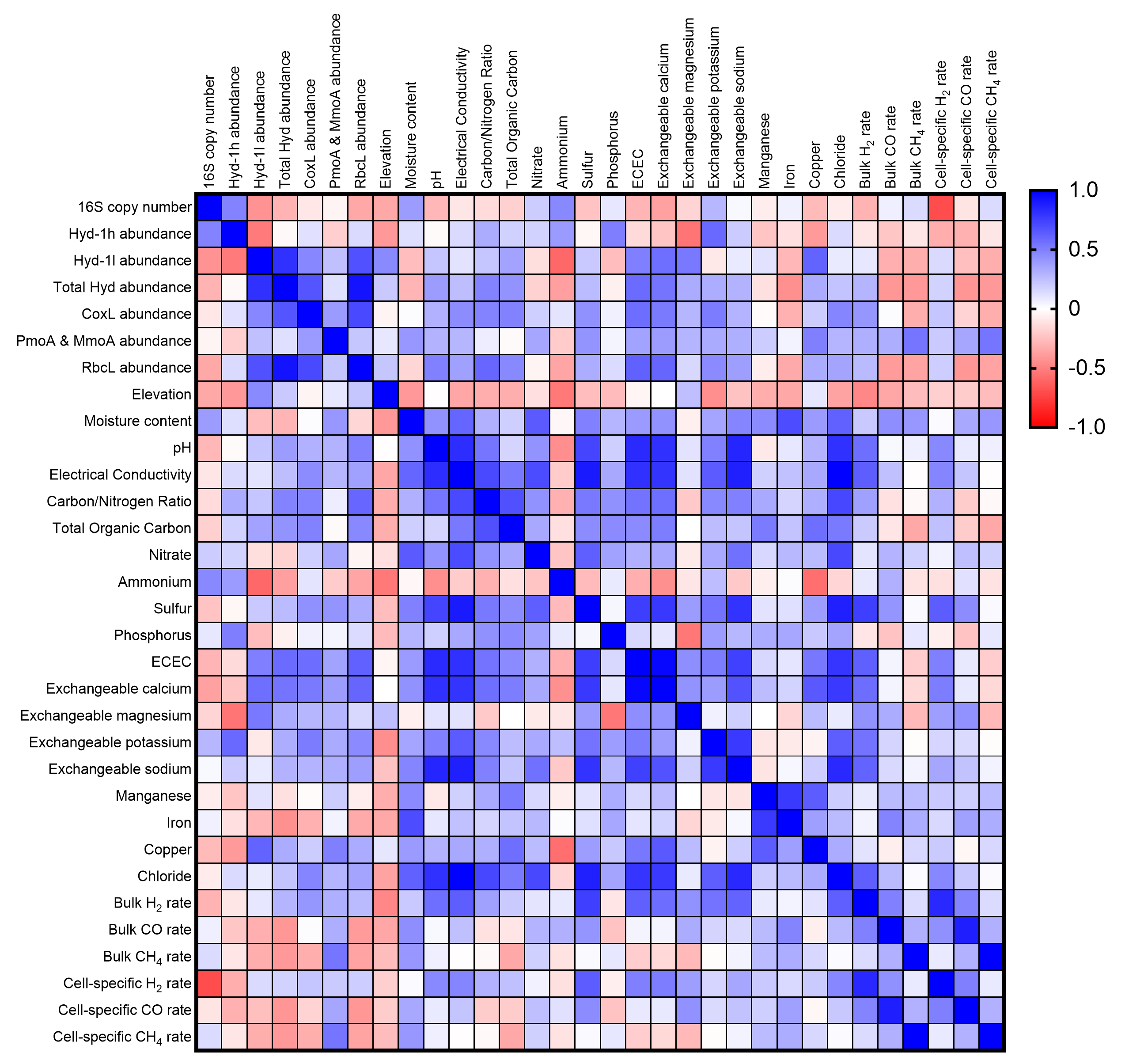

**Figure S7.** Gas chromatography measurement of hydrogen, carbon monoxide, and methane oxidation. Soils collected from the Mackay Glacier region were incubated over the timecourse at 10°C. Technical duplicate microcosms for each sampling site and quadruplicate microcosms for pooled heat-killed controls are shown with colored dots and solid lines. Dotted lines represent the mean atmospheric mixing ratios of the corresponding trace gas (H_2_: 0.53 ppmv; CO: 0.09 ppmv; CH_4_: 1.9 ppmv). Consumption to below atmospheric mixing ratios was observed for H_2_ (all soils except Pegtop Mountain), CO (10 out 16 sites), and CH_4_ (Mount Seuss 5) during the timecourse.

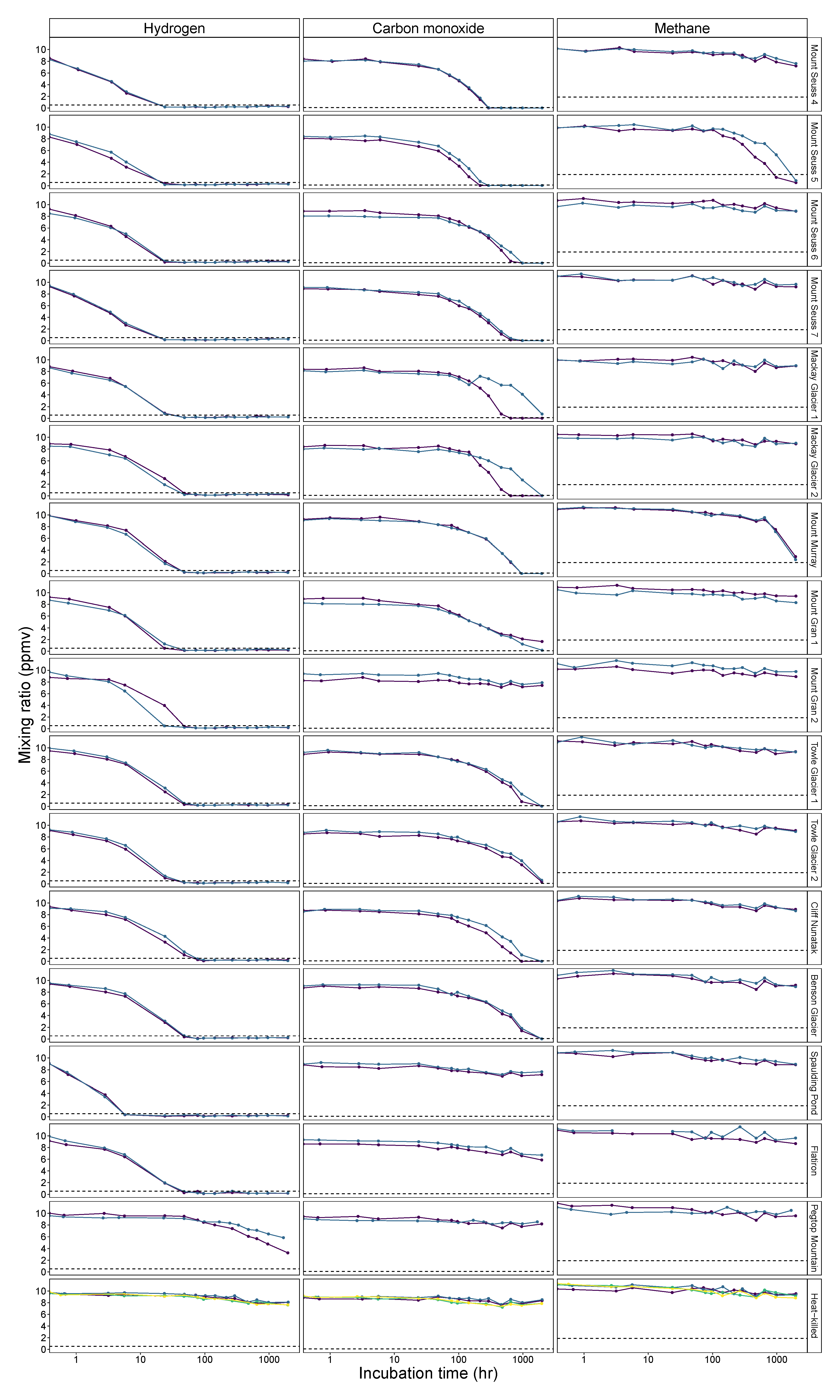

**Figure S8.** Maximum-likelihood tree of amino acid sequences of the ammonia monooxygenase A subunit (AmoA), a marker for aerobic ammonia oxidation during nitrification. The tree shows sequences from metagenome-assembled genomes (blue) and unbinned assembled sequences (red) from the Mackay Glacier region alongside representative reference sequences (grey). The tree was constructed using the JTT matrix-based model, used all sites, and was bootstrapped with 50 replicates and midpoint-rooted.

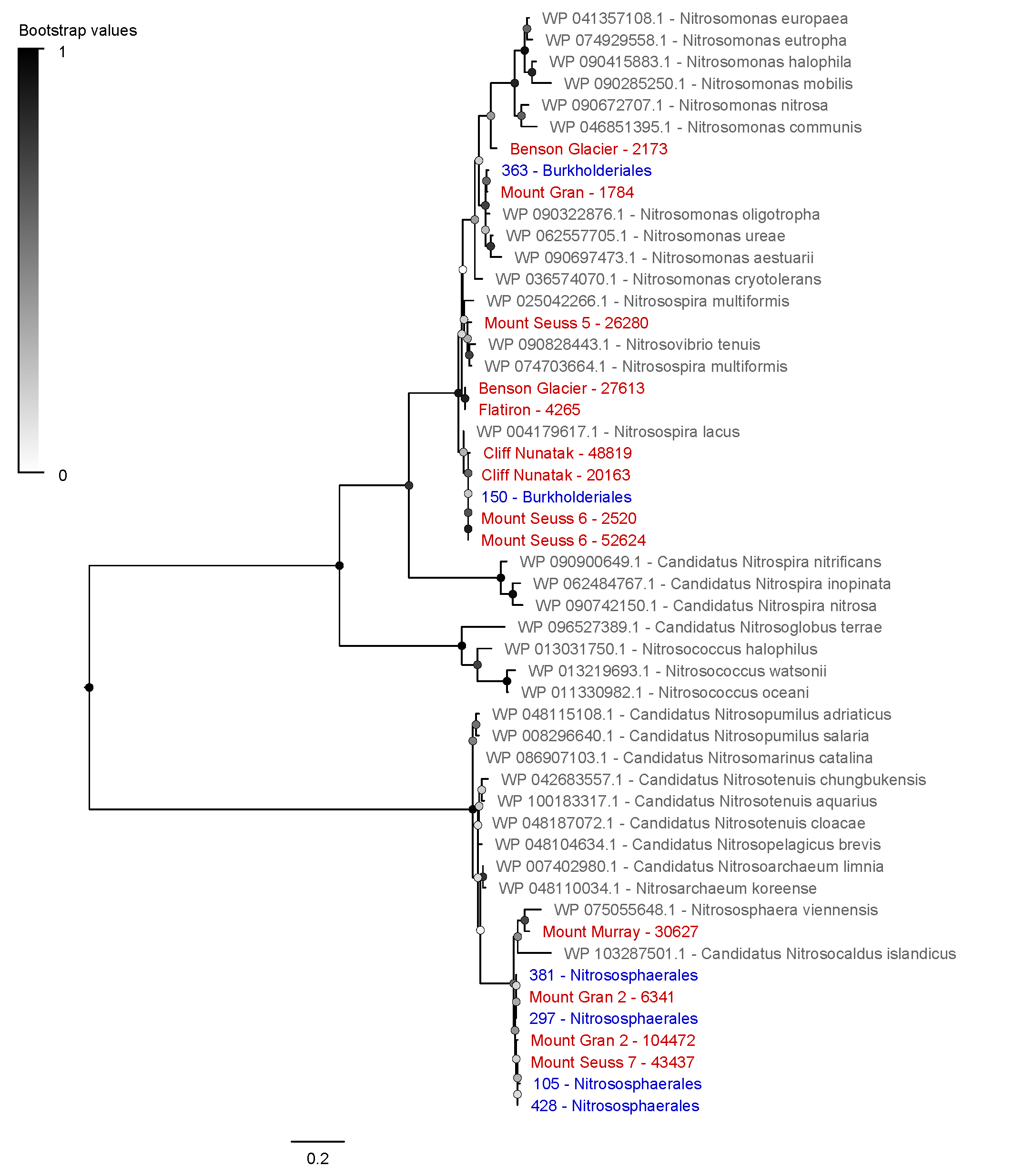

**Figure S9.** Maximum-likelihood tree of amino acid sequences of the nitrite oxidoreductase A subunit (NxrA), a marker for aerobic nitrite oxidation during nitrification. The tree shows sequences from metagenome-assembled genomes (blue) and unbinned assembled sequences (red) from the Mackay Glacier region alongside representative reference sequences (grey). The tree was constructed using the JTT matrix-based model, used all sites, and was bootstrapped with 50 replicates and midpoint-rooted.

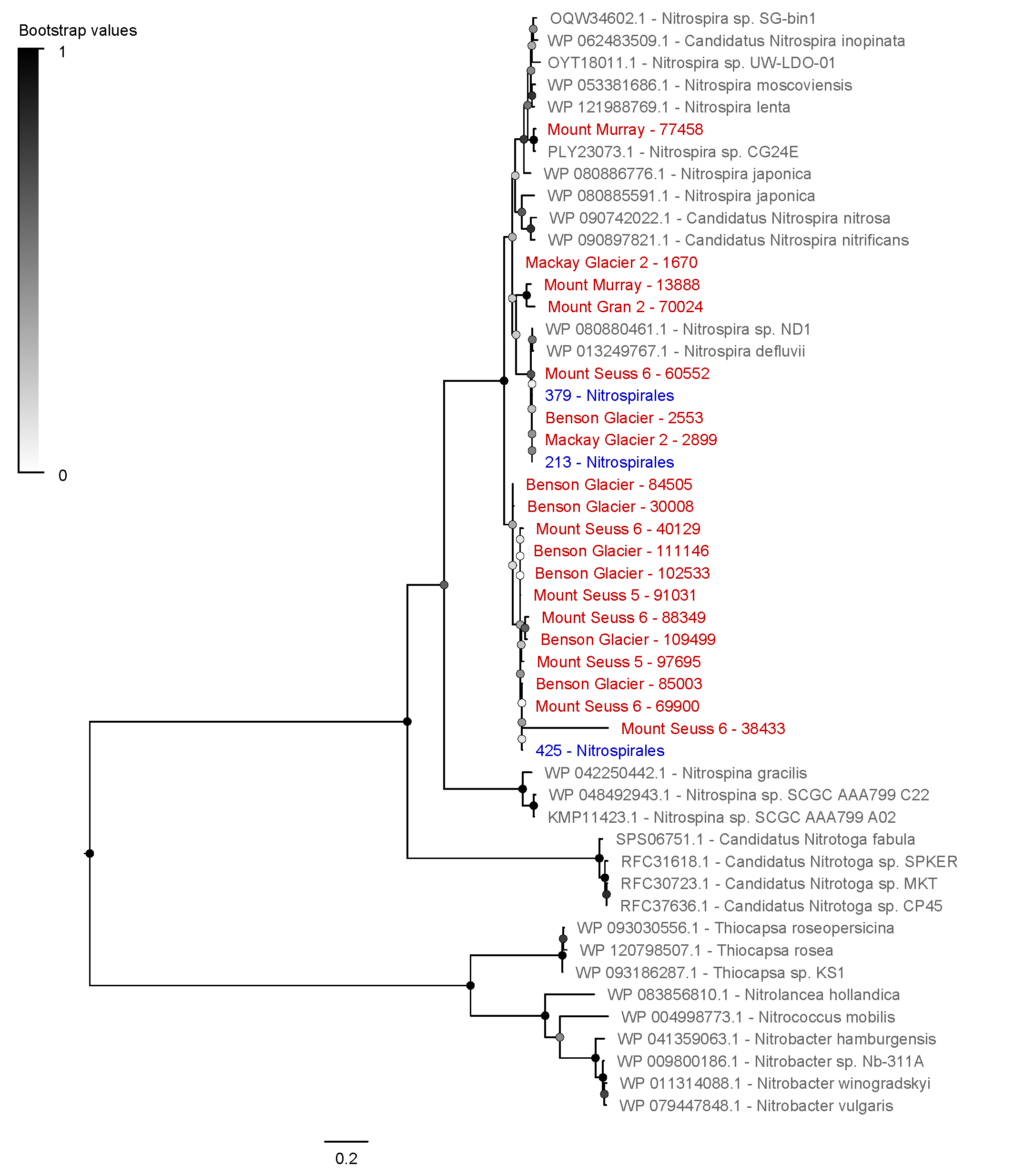

**Figure S10.** Maximum-likelihood tree of amino acid sequences of thaumarchaeotal 4-hydroxybutyryl-CoA synthetase (HbsT), a marker for carbon fixation through a variant of the 3-hydroxypropionate / 4-hydroxybutyrate cycle. The tree shows sequences from metagenome-assembled genomes (blue) and unbinned assembled sequences (red) from the Mackay Glacier region alongside representative reference sequences (grey). The tree was constructed using the JTT matrix-based model, used all sites, and was bootstrapped with 50 replicates and midpoint-rooted.

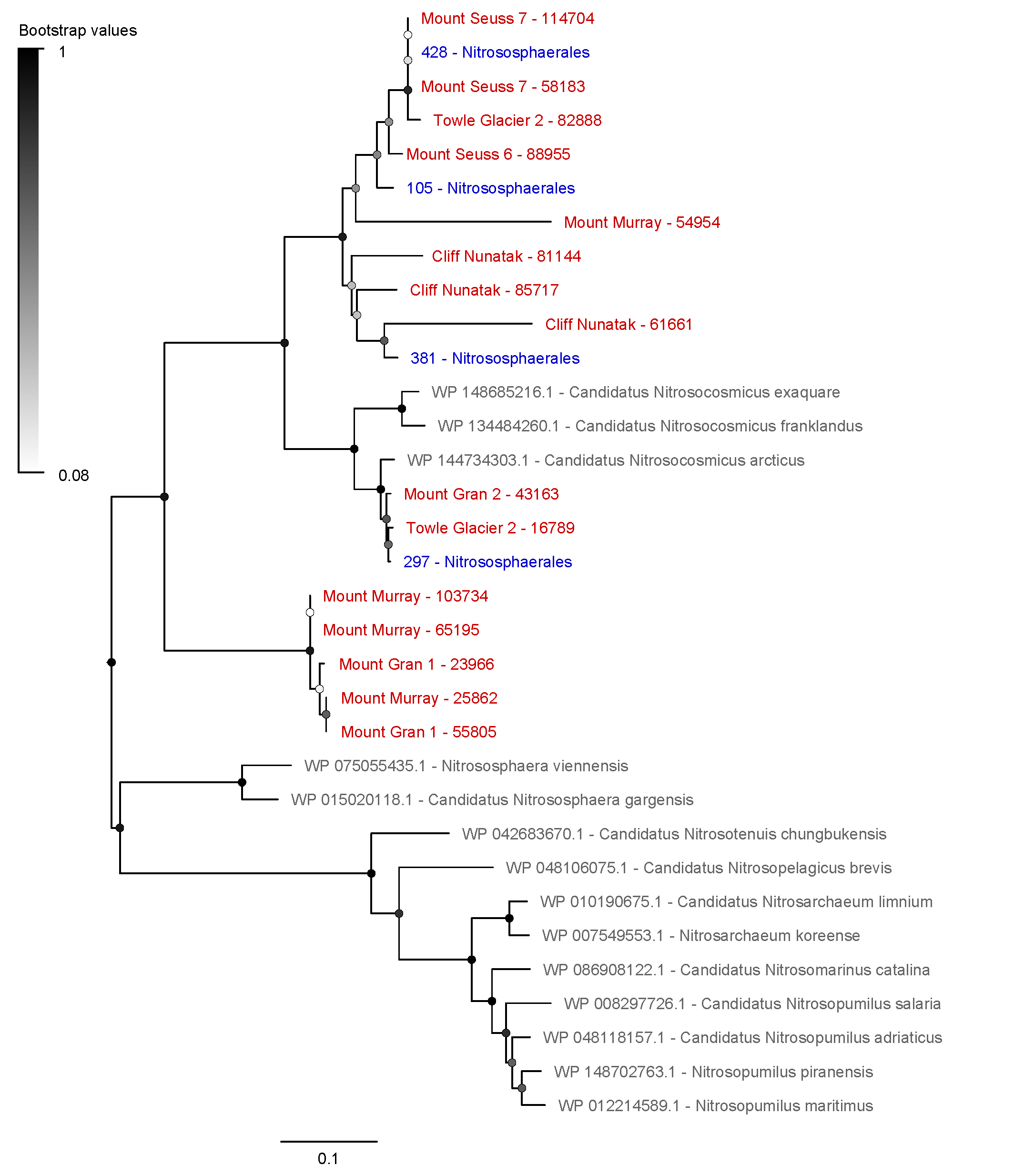

**Figure S11.** Maximum-likelihood tree of amino acid sequences of the ATP-citrate lyase B subunit (AclB), a marker for carbon fixation through the reductive tricarboxylic acid cycle. The tree shows sequences from metagenome-assembled genomes (blue) and unbinned assembled sequences (red) from the Mackay Glacier region alongside representative reference sequences (grey). The tree was constructed using the JTT matrix-based model, used all sites, and was bootstrapped with 50 replicates and midpoint-rooted.

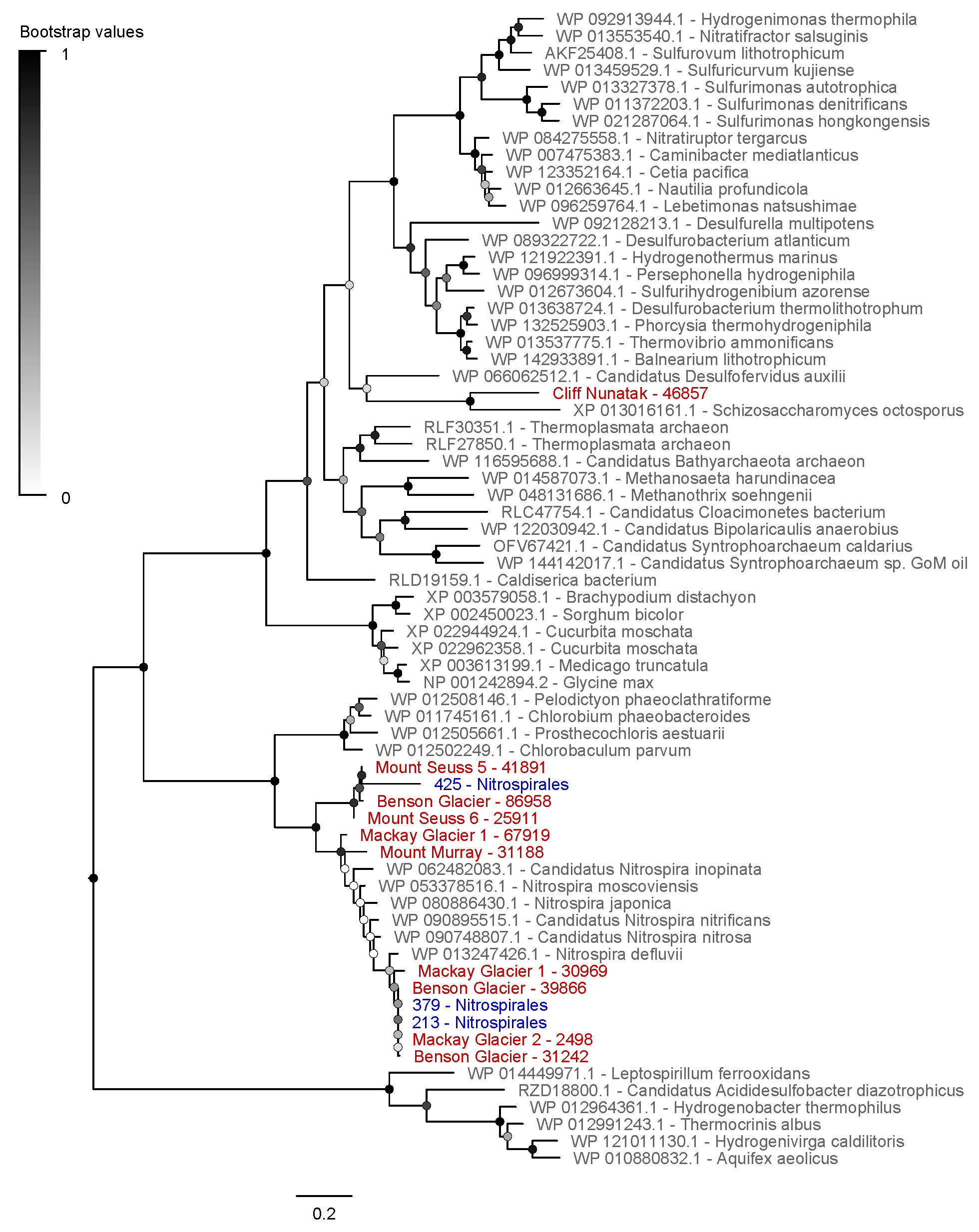

**Figure S12.** Maximum-likelihood tree of amino acid sequences of sulfide-quinone oxidoreductase (Sqr), a marker for aerobic sulfide oxidation. The tree shows sequences from metagenome-assembled genomes (blue) and unbinned assembled sequences (red) from the Mackay Glacier region alongside representative reference sequences (grey). The tree was constructed using the JTT matrix-based model, used all sites, and was bootstrapped with 50 replicates and midpoint-rooted.

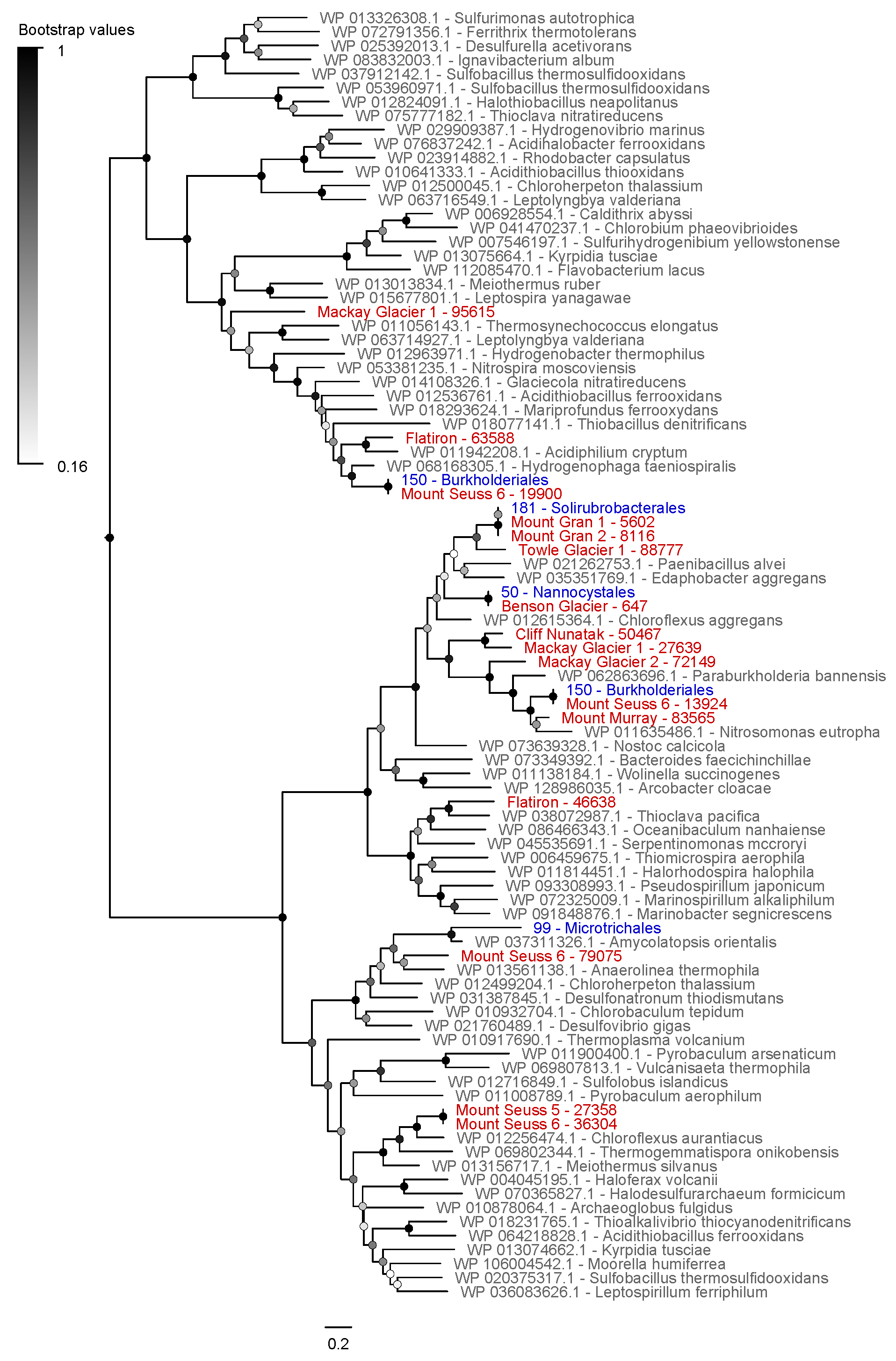

**Figure S13.** Maximum-likelihood tree of amino acid sequences of flavocytochrome *c* sulfide dehydrogenase (FCC), a marker for aerobic sulfide oxidation. The tree shows sequences from metagenome-assembled genomes (blue) and unbinned assembled sequences (red) from the Mackay Glacier region alongside representative reference sequences (grey). The tree was constructed using the JTT matrix-based model, used all sites, and was bootstrapped with 50 replicates and midpoint-rooted.

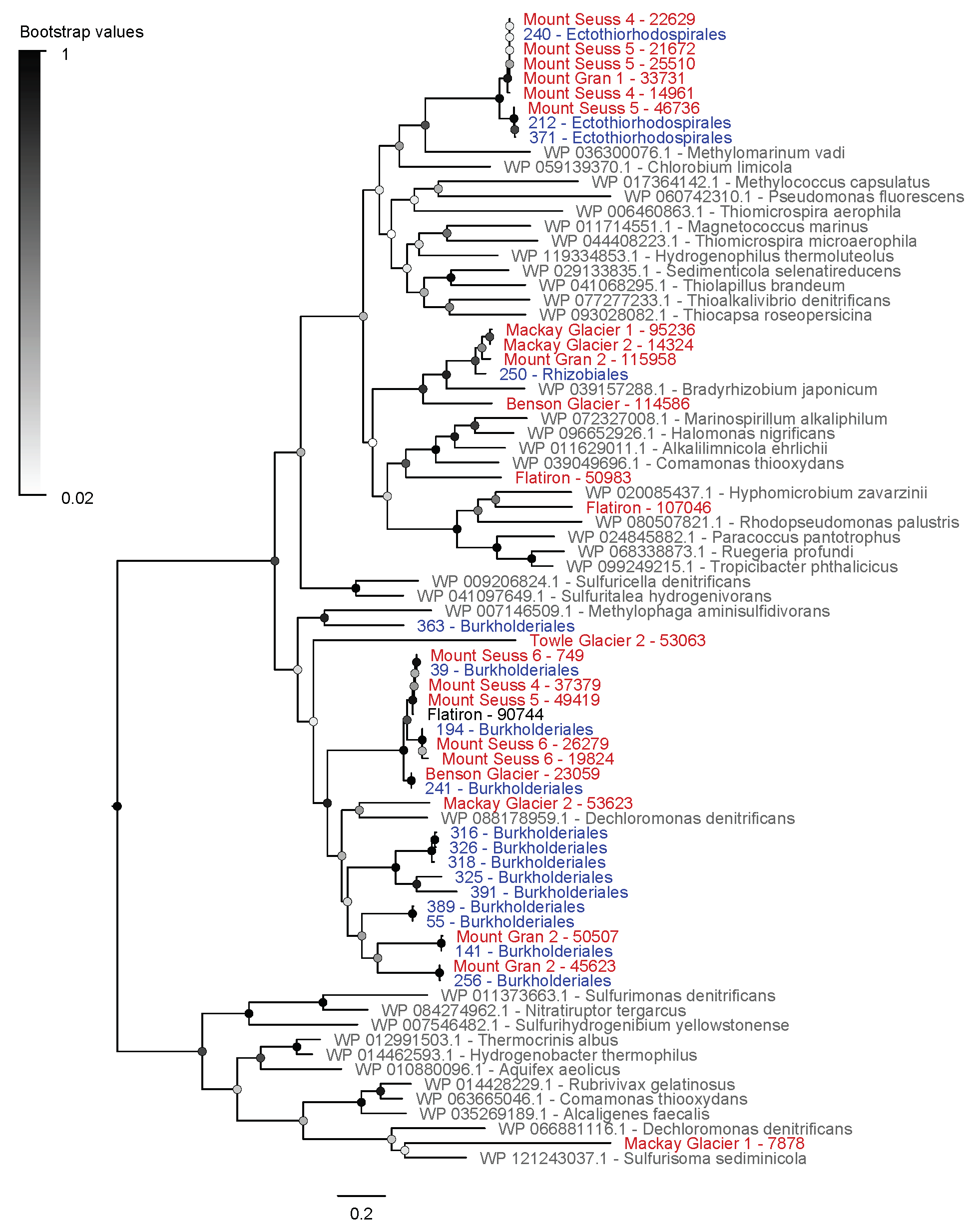

**Figure S14.** Maximum-likelihood tree of amino acid sequences of thiosulfohydrolase (SoxB), a marker for aerobic thiosulfate oxidation. The tree shows sequences from metagenome-assembled genomes (blue) and unbinned assembled sequences (red) from the Mackay Glacier region alongside representative reference sequences (grey). The tree was constructed using the JTT matrix-based model, used all sites, and was bootstrapped with 50 replicates and midpoint-rooted.

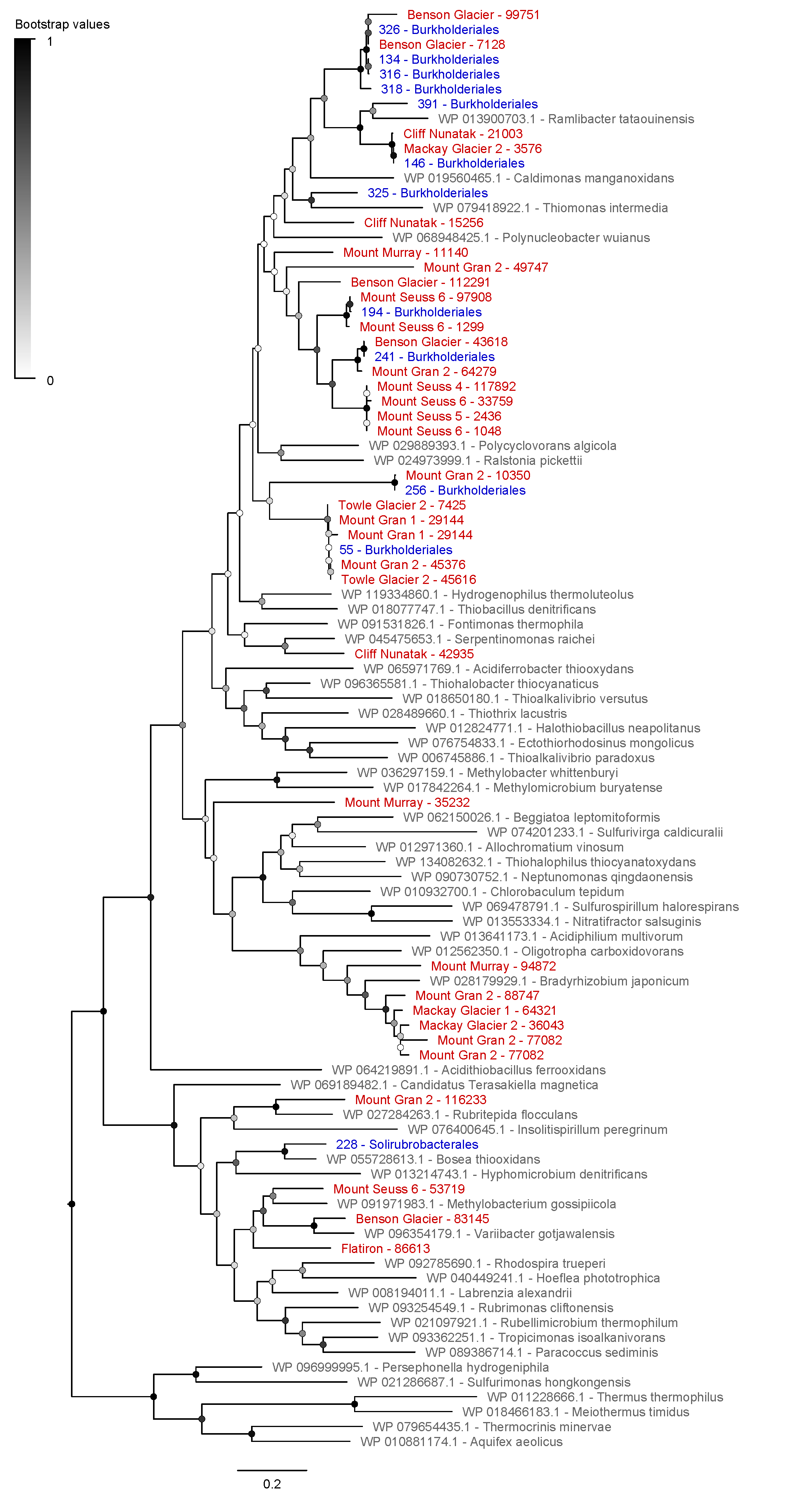

**Figure S15.** Maximum-likelihood tree of amino acid sequences of iron-oxidizing *c*-type cytochrome (Cyc2), a marker for aerobic Fe(II) oxidation. The tree shows sequences from metagenome-assembled genomes (blue) and unbinned assembled sequences (red) from the Mackay Glacier region alongside representative reference sequences (grey). The tree was constructed using the JTT matrix-based model, used all sites, and was bootstrapped with 50 replicates and midpoint-rooted.

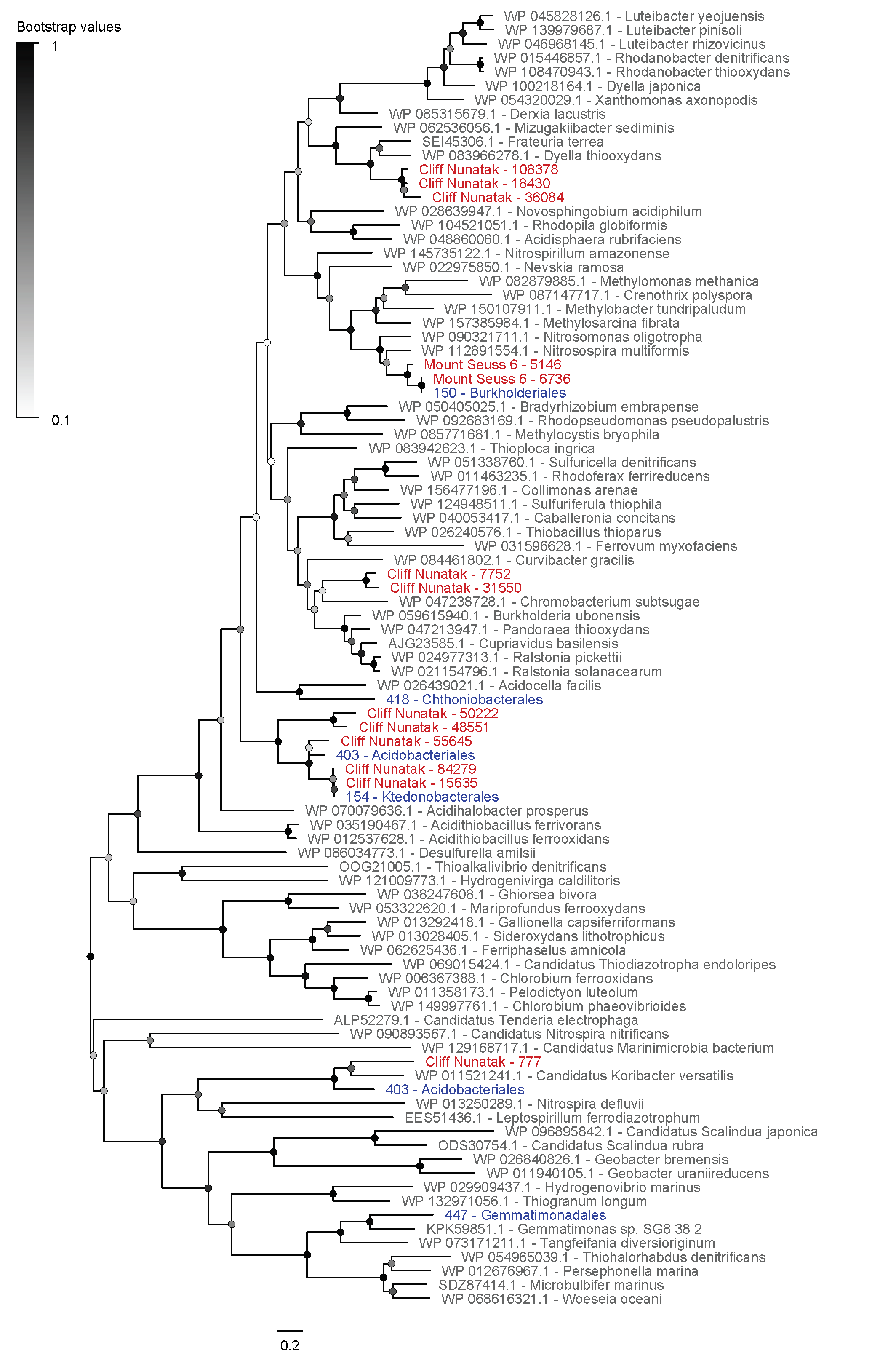

**Figure S16.** Maximum-likelihood tree of amino acid sequences of the photosystem II A subunit (PsbA), a marker for oxygenic and anoxygenic photosynthesis. The tree shows sequences from metagenome-assembled genomes (blue) and unbinned assembled sequences (red) from the Mackay Glacier region alongside representative reference sequences (grey). The tree was constructed using the JTT matrix-based model, used all sites, and was bootstrapped with 50 replicates and midpoint-rooted.

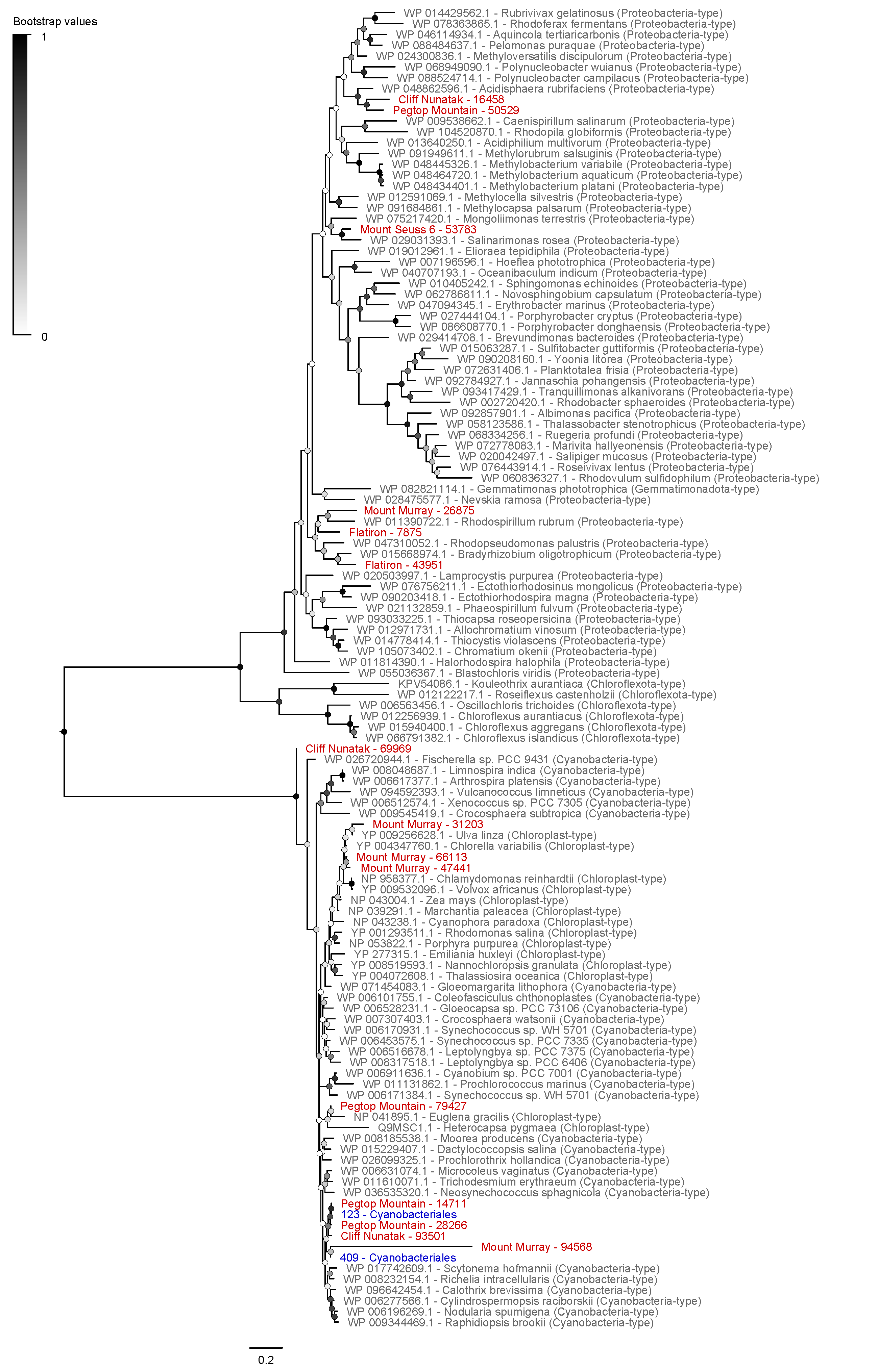

**Figure S17.** Maximum-likelihood tree of amino acid sequences of energy-converting rhodopsins. The tree shows sequences from metagenome-assembled genomes (blue) and unbinned assembled sequences (red) from the Mackay Glacier region alongside representative reference sequences (grey). Two clades are likely to be novel families of energy-converting rhodopsins. The tree was constructed using the JTT matrix-based model, used all sites, and was bootstrapped with 50 replicates and midpoint-rooted.

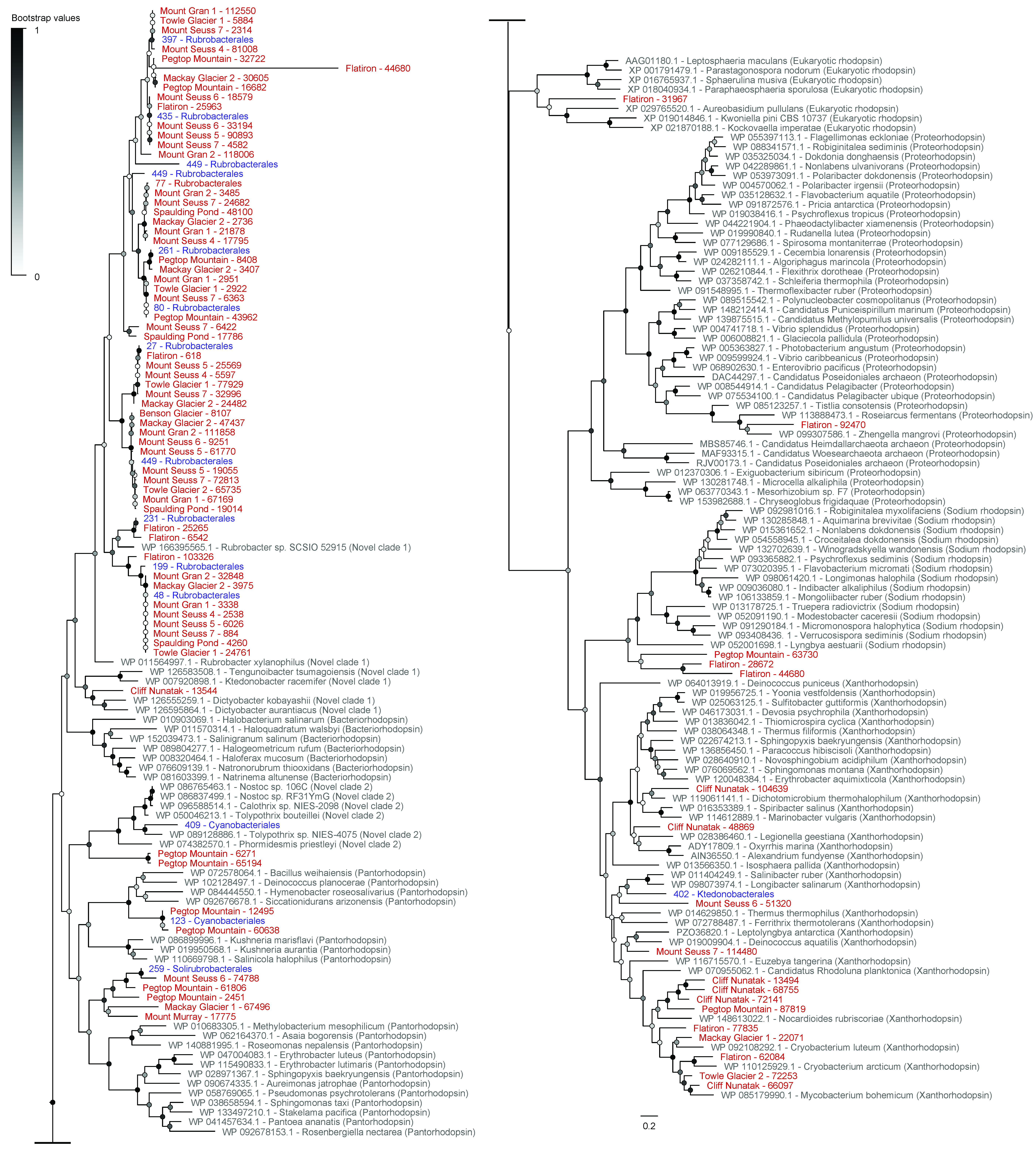

**Figure S18.** Maximum-likelihood tree of amino acid sequences of the particulate methane monooxygenase A subunit (PmoA), a marker for aerobic methane oxidation. The tree shows sequences from metagenome-assembled genomes (blue) and unbinned assembled sequences (red) from the Mackay Glacier region alongside representative reference sequences (grey). The tree was constructed using the JTT matrix-based model, used all sites, and was bootstrapped with 50 replicates and midpoint-rooted.

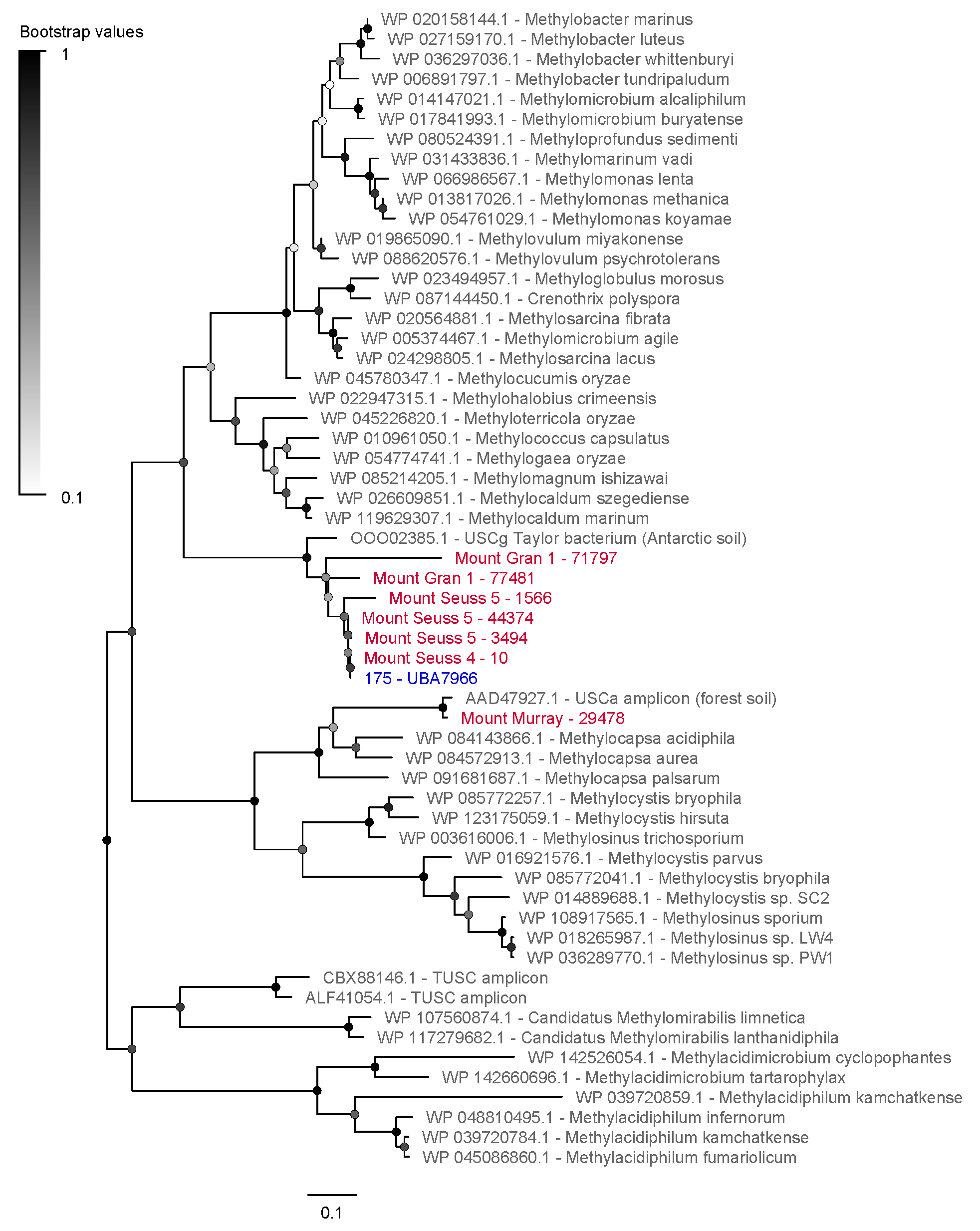
